# Conversion of monoclonal IgG to dimeric and secretory IgA restores neutralizing ability and prevents infection of Omicron lineages

**DOI:** 10.1101/2023.04.17.536908

**Authors:** Harold Marcotte, Yunlong Cao, Fanglei Zuo, Luca Simonelli, Josè Camilla Sammartino, Mattia Pedotti, Rui Sun, Irene Cassaniti, Marie Hagbom, Antonio Piralla, Jinxuan Yang, Likun Du, Elena Percivalle, Federico Bertoglio, Maren Schubert, Hassan Abolhassani, Natalia Sherina, Concetta Guerra, Stephan Borte, Nima Razaei, Makiko Kumagai-Braesch, Yintong Xue, Caroline Grönwall, Lars Klareskog, Luigi Calzolai, Andrea Cavalli, Qiao Wang, Davide F. Robbiani, Michael Hust, Zhengli Shi, Liqiang Feng, Lennart Svensson, Ling Chen, Linlin Bao, Fausto Baldanti, Chuan Qin, Junyu Xiao, Lennart Hammarström, Xing Lou Yang, Luca Varani, Xiaoliang Sunney Xie, Qiang Pan-Hammarström

**Affiliations:** Department of Biosciences and Nutrition, Karolinska Institutet, Huddinge, Sweden; Changping Laboratory, Beijing, P.R. China; Biomedical Pioneering Innovation Center (BIOPIC), Peking University, Beijing, P. R. China; Institute for Research in Biomedicine, Università della Svizzera Italiana (USI), Bellinzona, Switzerland; Microbiology and Virology Department, Fondazione IRCCS Policlinico San Matteo, Pavia, Italy; Division of Molecular Medicine and Virology, Linköping University, Linköping, Sweden; Kunming Institute of Zoology, CAS, Panlong district, Kunming city, China; Technische Universität Braunschweig, Institute of Biochemistry, Biotechnology and Bioinformatics, Department of Medical Biotechnology, Braunschweig, Germany; Department of Laboratory Medicine, Hospital St. Georg, Leipzig, Germany; ImmunoDeficiencyCenter Leipzig, Jeffrey Modell Diagnostic and Research Center for Primary Immunodeficiency Diseases, Hospital St. Georg, Leipzig, Germany; Research Center for Immunodeficiencies, Pediatrics Center of Excellence, Children’s Medical Center, Tehran University of Medical Sciences, Tehran, Iran; Division of Transplantation Surgery, CLINTEC, Karolinska Institutet at Karolinska University Hospital, Stockholm, Sweden; Department of Immunology, Peking University Health Science Center, Beijing, China; Department of Medicine Solna, Division of Rheumatology, Center for Molecular Medicine, Karolinska Institutet, Karolinska University Hospital, Stockholm, Sweden; Rheumatology Clinic, Karolinska University Hospital, Stockholm, Sweden; European Commission, Joint Research Centre, Ispra, Italy; Key Laboratory of Medical Molecular Virology (MOE/NHC/CAMS), Shanghai Institute of Infectious Disease and Biosecurity, School of Basic Medical Sciences, Fudan University, Shanghai, China; CAS Key Laboratory of Special Pathogens, Wuhan Institute of Virology, Chinese Academy of Sciences, Wuhan, P.R. China; Bioland Laboratory (GRMH-GDL), Guangzhou Institutes of Biomedicine and Health, Chinese Academy of Sciences, Guangzhou, China; State Key Laboratory of Respiratory Disease, Guangzhou Institute of Respiratory Health, First Affiliated Hospital of Guangzhou Medical University, Guangzhou, China; Division of Infectious Diseases, Department of Medicine, Karolinska Institute, Solna, Sweden; Beijing Key Laboratory for Animal Models of Emerging and Remerging Infectious Diseases, NHC Key Laboratory of Human Disease Comparative Medicine, Institute of Laboratory Animal Science, Chinese Academy of Medical Sciences and Comparative Medicine Center, Peking Union Medical College, Beijing, China; National Center of Technology Innovation for Animal Model, Beijing, China; Department of Clinical, Surgical, Diagnostic and Paediatric Sciences, University of Pavia, Pavia, Italy

## Abstract

The emergence of Omicron lineages and descendent subvariants continues to present a severe threat to the effectiveness of vaccines and therapeutic antibodies. We have previously suggested that an insufficient mucosal IgA response induced by the mRNA vaccines is associated with a surge in breakthrough infections. Here, we further show that the intramuscular mRNA and/or inactivated vaccines cannot sufficiently boost the mucosal sIgA response in uninfected individuals, particularly against the Omicron variant. We thus engineered and characterized recombinant monomeric, dimeric and secretory IgA1 antibodies derived from four neutralizing IgG monoclonal antibodies targeting the receptor-binding domain of the spike protein (01A05, rmAb23, DXP-604 and XG014). Compared to their parental IgG antibodies, dimeric and secretory IgA1 antibodies showed a higher neutralizing activity against different variants of concern (VOCs), in part due to an increased avidity. Importantly, the dimeric or secretory IgA1 form of the DXP-604 antibody significantly outperformed its parental IgG antibody, and neutralized the Omicron lineages BA.1, BA.2 and BA.4/5 with a 50-150-fold increase in potency, reaching the level of the most potent monoclonal antibodies described till date. In hACE2 transgenic mice, a single intranasal dose of the dimeric IgA DXP-604 conferred prophylactic and therapeutic protection against Omicron BA.5. Conversion of IgA and dimerization further enhanced or restored the neutralizing ability against the emerging Omicron sub-variants (DXP-604 for BQ.1, BQ.1.1 and BA2.75; 01A05 for BA2.75, BA.2.75.2 and XBB.1). Thus, dimeric or secretory IgA delivered by nasal administration may potentially be exploited for the treatment and prevention of Omicron infection, thereby providing an alternative tool for combating immune evasion by subvariants and, potentially, future VOCs.

**One Sentence Summary:** Engineered dimeric and secretory IgA1 neutralized Omicron variant with higher potency than parental IgG.

## INTRODUCTION

As severe acute respiratory syndrome coronavirus 2 (SARS-CoV-2) continues to spread worldwide, selection pressure to evade antibodies in convalescent and/or vaccinated individuals has led to viral mutations and the emergence of Variants of Concerns (VOCs), such as Alpha (B.1.1.7), Beta (B.1.351), Gamma (P.1) and Delta (B.1.617.2) *(1)*. Mutations in the viral spike (S) protein, including the receptor-binding domain (RBD), may lead to a reduced susceptibility to neutralization by antibodies, an increased binding to the ACE2 receptor on host cells and a higher transmissibility and infectivity *(1)*. The emergence of the Omicron variant in South Africa in November 2021, and its rapid spread worldwide have strengthened concerns about vaccine efficacy and antibody therapy due to the large number of mutations in the S protein *(2–5)*. The original Omicron variant (B.1.1.529) already harbours 37 mutations in the S glycoprotein, including 15 in the RBD *(6)*. It is continuously evolving, and has hitherto divided into five major lineages, BA.1 to BA.5 *(7)*. Novel subvariants such as BA.2.75, BQ.1, and XBB, derived from either BA.2 or BA.4/5, have rapidly emerged *(8)*. The rise of these subvariants seems to stem from their capacity to evade the immune system and infect individuals who are immune to earlier Omicron subvariants *(9, 10)*.

The development of novel antibody therapies that remain efficacious despite virus evolution is thus urgently needed. Widespread reinfections and vaccine breakthrough infections (BTIs) with Omicron have been reported worldwide, and most of the clinically available antibodies have been found to be ineffective against this variant *(5, 11, 12)*. Although Omicron causes less severe symptoms than previous variants, it still results in a substantial number of hospitalizations and deaths, especially in unvaccinated individuals. Antibody pre-exposure therapy could be beneficial for the protection of individuals at high risk of developing severe diseases, such as immunocompromised patients and elderly individuals, especially in areas/countries with low vaccination/booster rates *(13–16)*. Recent evidence suggests a shift in the tropism of the Omicron variant towards the upper respiratory tract *(17)*. Viral particles in the upper airways might be more easily released from the nose and mouth, contributing to the increased transmissibility of the Omicron variant *(18)*. The virus might be contained in the upper respiratory tract of individuals who develop a strong local mucosal immune response, resulting in a mild/asymptomatic infection *(19)*. Thus, mucosal immunity may potentially be exploited for therapeutic or prophylactic purposes *(20)*.

Secretory IgA (sIgA) is the most abundant immunoglobulin type in secretions and is fundamental for mucosal defences and protection against respiratory viral infections. While serum IgA is predominantly present as a monomer (mIgA), sIgA is composed of two IgA monomers, connected via the joining (J) chain, and associated with the secretory component (SC) *(21)*. Dimeric IgA (dIgA) produced by B cells in the mucosa is translocated across the epithelium via the polymeric immunoglobulin receptor (pIgR) *(22)*. On the luminal side of the epithelium, pIgR is cleaved, while a portion, the SC, remains attached, forming sIgA *(23)*. Among the two subclasses of IgA antibodies in humans, IgA1 constitutes a higher proportion in the upper respiratory tract (approximately 90%), and IgA2 is more abundant in the lower gastrointestinal tract *(23–25)*. Mucosal IgA dominates the neutralizing antibody response to SARS-CoV-2 in the early phase of infection and is more effective in neutralizing SARS-CoV-2 *(26, 27)*, and dIgA has been found to be a more potent neutralizer than IgG against authentic SARS-CoV-2 *(28)*. Thus, delivery of both dIgA and sIgA via nasal spray is potentially the best and most convenient option to block viral infection and transmission.

## RESULTS

### Secretory anti-RBD IgA antibodies are produced at low levels following vaccination

Among immunoglobulins, sIgA are present at highest concentration in saliva and constitute an accessible marker of the mucosal immune response to SARS-CoV-2 *(29, 30)*. We previously showed that in individuals receiving mRNA vaccines, higher levels of salivary anti-RBD sIgA antibodies were associated with protection against BTI *(31)*. Here, we further tested the presence of salivary antibodies against the RBDs of G614 (wild-type) and of all Omicron lineages in individuals who received various doses of either inactivated or mRNA vaccines (**Fig. 1, A to C and table S1**) *(31)*.

**Fig. 1.**
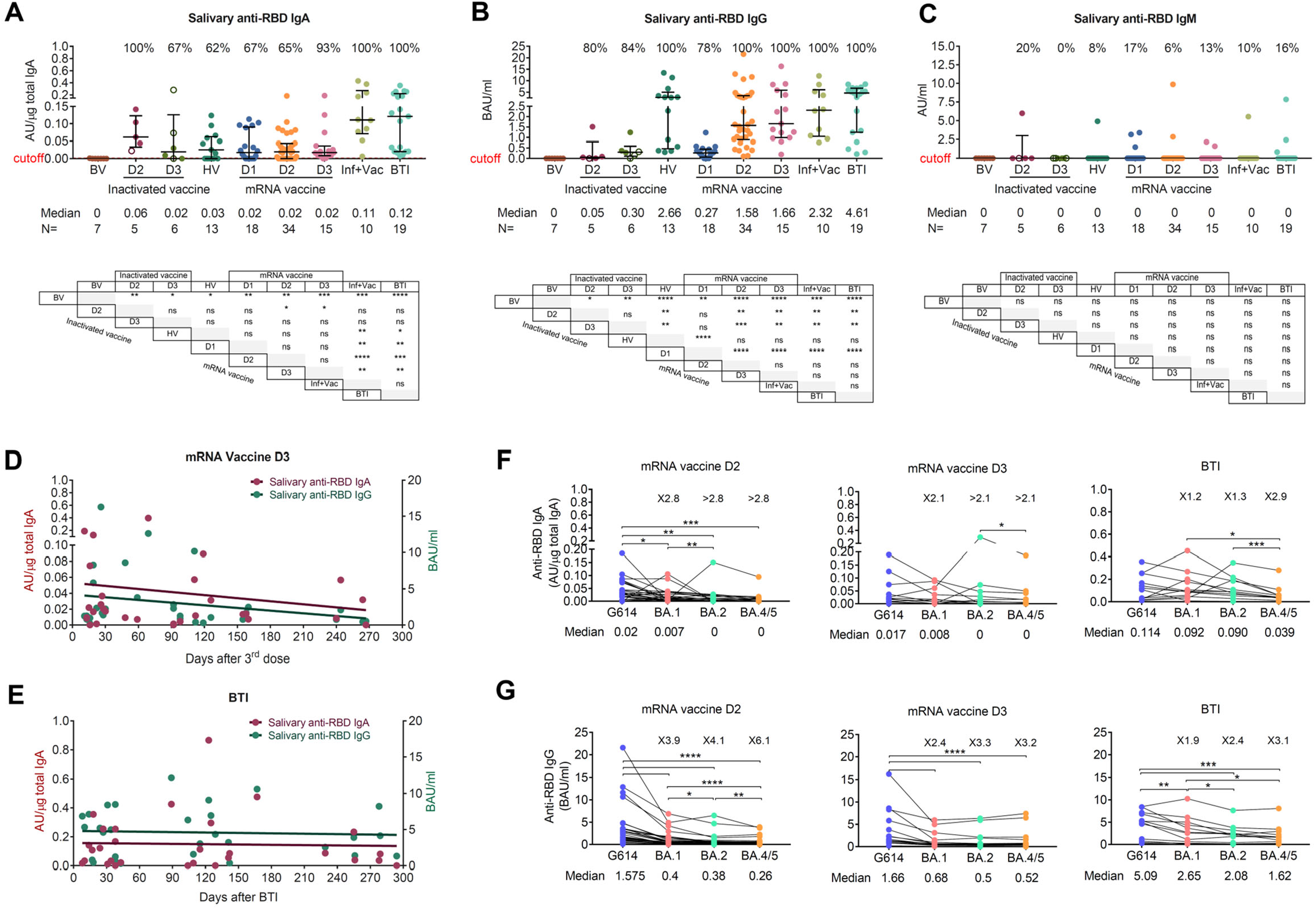
Salivary anti-RBD IgA antibodies are produced at low levels following vaccination. (**A** to **C**) Salivary anti-RBD IgA (A), IgG (B) and IgM (C) antibodies in different vaccination groups. For each group, the number of samples (n =) and median antibody titres are shown below the X-axis. Whiskers indicate the interquartile range. The results of anti-RBD antibodies are presented as arbitrary units (AU)/µg total IgA (salivary IgA), binding antibody units (BAU/ml) (salivary and plasma IgG) or arbitrary units (AU)/ml (salivary IgM). HV: Heterologous vaccination (two doses of inactivated vaccine followed by a heterologous mRNA boost, Inf+Vac: one or two doses of mRNA vaccine after SARS-CoV-2 infection (during the G614 wave), BTI: breakthrough infection (during the BA.1 wave) after inactivated and/or mRNA vaccines. A two-sided Mann–Whitney U test was used. (**D** and **E**) Dynamics of salivary anti-RBD IgA and IgG antibodies after the third (D3) doses of mRNA vaccine (BNT162b2 or mRNA-1273) (D) or after a breakthrough infection (BTI) in mRNA-vaccinated individuals (E). (**F** and **G**) Salivary anti-RBD IgA (F) and IgG (G) antibodies against G614 and Omicron variants BA.1, BA.2 and BA.4/5 after the second (D2) and third (D3) doses of mRNA vaccine, and following BTI in mRNA-vaccinated individuals. The number of fold differences of median compared to G614 are indicated. A Wilcoxon paired-sample signed-rank test was used. *p < 0.05, ** p < 0.01, *** p < 0.001, and **** p < 0.0001. ns, not significant. See also fig. S1.

The results show that the median salivary anti-RBD IgA levels in individuals receiving two or three doses of inactivated whole-virion SARS-CoV-2, one to three doses of mRNA vaccine or heterologous vaccination were lower than those after BTI or an mRNA vaccine booster after infection (**Fig. 1A**). Salivary anti-RBD IgG levels after the second and third doses of mRNA vaccine or heterologous mRNA booster dose were similar to those measured after BTI (**Fig. 1B)** while salivary IgM anti-RBD antibodies were detected in less than 20% of individuals after vaccination and BTI (**Fig. 1C)**. Salivary RBD-specific IgA antibodies gradually decreased after the third mRNA vaccine dose but were still detectable in nearly 85% of individuals up to 9 months, while after BTI, these antibodies were sustained at a higher level (**Fig. 1, D and E**). Lower salivary IgA (from 2.1- to > 2.8-fold) and IgG (from 2.4- to 6.1-fold) antibody levels against the RBDs of BA.1, BA.2 and BA.4/5 compared to G614 RBD were observed in vaccinated individuals within two months after two or three doses of mRNA vaccine, but no decrease in antibody levels against Omicron variants was observed in individuals who experienced BTIs during the Omicron BA.1 wave, suggesting a long-lasting and broadly cross-reactive mucosal immune response after BTI (**Fig. 1, F and G**). Furthermore, a significant increase (by more than 10-fold) in salivary IgA and IgG antibodies against Omicron subvariants RBD was observed 2-6 weeks after BTI (**fig. S1, A and B**).

Levels of salivary anti-RBD IgA antibodies correlated better with levels of RBD-specific secretory immunoglobulin (sIg) (R = 0.789, p *<* 0.0001) than plasma IgA antibodies (R = 0.256, p *<* 0.0046) (**fig. S1C and D**), confirming that most of the salivary IgA antibodies measured were produced locally in the salivary glands as sIgA *(31)*. Salivary anti-RBD IgG levels correlated with plasma IgG antibodies, implying that those antibodies were mainly derived from plasma (R = 0.704, p *<* 0.0001) (**fig. S1E**).

In conclusion, in uninfected individuals, vaccination cannot efficiently induce/boost the mucosal IgA response against SARS-CoV-2, especially the Omicron lineages. Furthermore, we and others have previously suggested that insufficient mucosal IgA response is associated with breakthrough infections during the Omicron waves *(27, 31)*. The development of vaccination strategies that can improve the mucosal IgA response, together with an effective IgA monoclonal antibody therapy that can be delivered directly at the mucosal surface, are therefore priorities in the current stage of the pandemic.

### Characterization of parental neutralizing IgG antibodies

Four neutralizing IgG mAbs, 01A05 (isolated in this study), rmAb23 *(32)*, DXP-604 *(2, 33)*, and XG014 *(34, 35)*, targeting SARS-CoV-2 RBD, were used for conversion into IgA formats. All IgG mAbs were originally isolated from convalescent donors who had the wild-type strain (Wuhan or G614 strain) infection before the emergence of Omicron *(32–35)*. Neutralizing anti-SARS-CoV-2 antibodies can be categorized into four *(36)* classes based on their mode of binding to the S protein. Based on computational analysis and structure, 01A05, encoded by IGHV1-18 and IGKV1-39, and rmAb23 and DXP-604, both encoded by IGHV3-53 and IGKV1-9 *(32)*, were shown to be class I antibodies that bind to the RBD in the up conformation (**Fig. 2A and fig. S2, A to C**). XG014 (IGHV5-51/IGLV1-51) is a class IV antibody that recognizes a conserved epitope outside the receptor-binding motif (RBM) in the RBD and locks all three RBDs in the S-trimer in the down conformation, preventing binding to ACE2 *(34)* (**Fig. 2A and fig. S2D**). In summary, the four antibodies recognize different epitopes in the RBD, with the Fabs of 01A05 and rmAb23 binding G614 RBD with a lower affinity (a dissociation constant (K_D_) of 2.5 nM and 6.5 nM *(32)*, respectively) compared to those of DXP-604 and XG014, which show subnanomolar K_D_ *(2, 33–35)*.

**Fig. 2.**
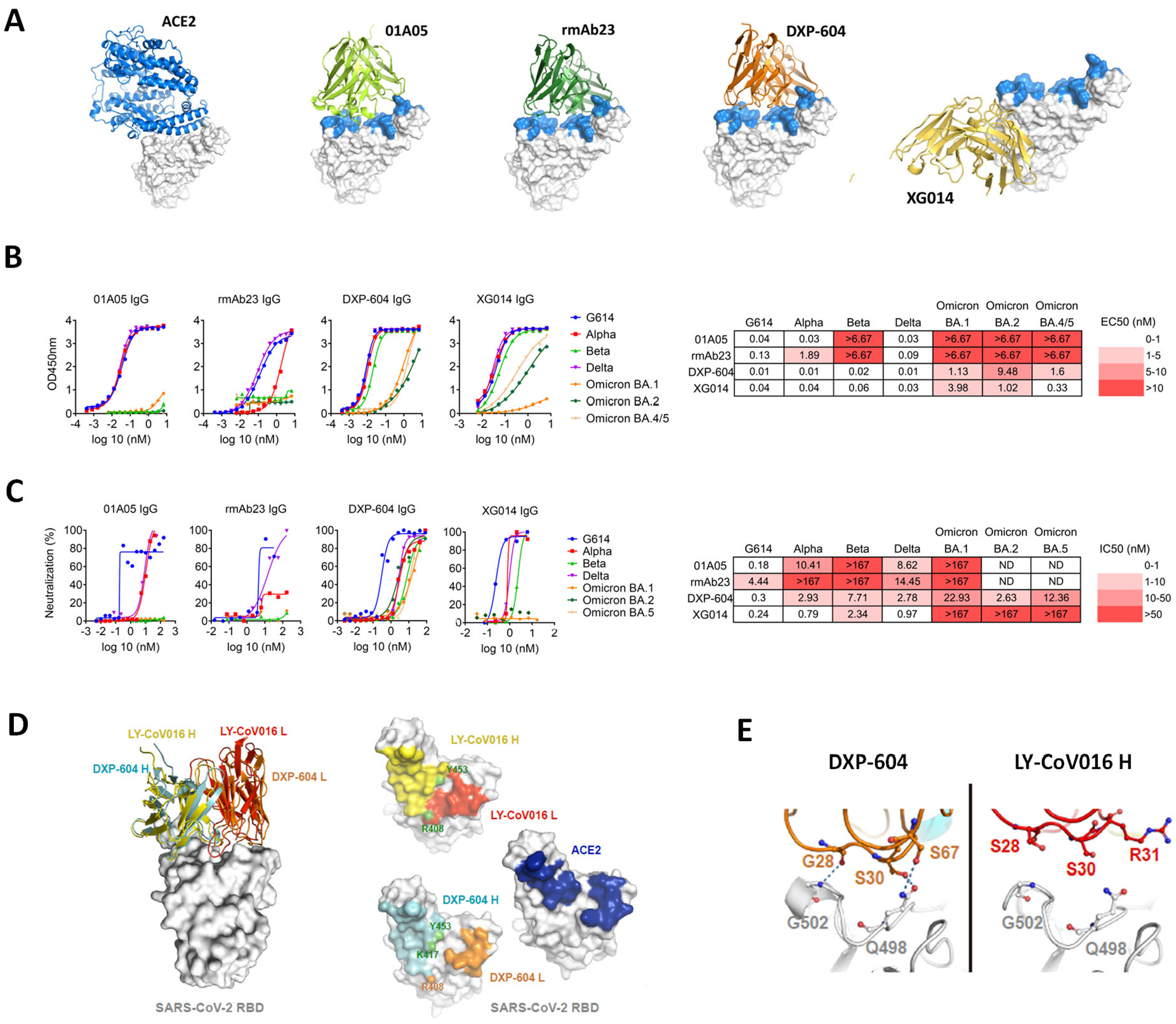
Characterization of neutralizing antibodies 01A05, rmAb23, DXP-604 and XG014. (**A**) In silico binding of ACE2 and IgG antibodies to RBD. The ACE2 receptor binding motif (RBM) is indicated (blue). (**B** and **C**) Binding to RBD (B) and neutralization (C) of G614 and variants of concern (VOCs) by IgG antibodies. The EC_50_ and IC_50_ and fold-change differences between IgG and IgA antibody forms are indicated. (**D**) Overlaid crystal structures of LY-CoV016 Fab (PDB ID: 7C01) and DXP-604 Fab 473 (PDB ID: 7CH4) in complex with SARS-CoV-2 RBD (left picture) and the footprints of LY-CoV016, DXP-604, and ACE2 (PDB ID: 6M0J) on SARS-CoV-2 RBD. Atoms of the RBD within 5.0 Å of the antibodies or ACE2 are coloured yellow (LY-CoV016 H), red (LY-CoV016 L), cyan (DXP-604 H), orange (DXP-604 L) or blue (ACE2) (right picture). (**E**) Hydrogen bonds were formed between S30/S67 in the light chain of DXP-604 and RBD Q498, which is a key ACE2-binding site, and between the main chain groups of Q27/S28 and RBD G502. See also fig. S2, S3 and S4.

All four antibodies bound RBDs from G614, Alpha and Delta with a one-half maximal effective concentration (EC_50_) value from 0.01 to 1.89 nM **(Fig. 2B)** but only DXP-604 and XG014 bound to RBDs from Beta, and Omicron BA.1, BA.2, BA.4/5 (EC_50_ from 0.33 to 9.48 nM) (**Fig. 2B**). In accordance with the ELISA results, 01A5 neutralized only G614, Alpha and Delta (half maximal inhibitory concentration (IC_50_): 0.18, 10.41 and 8.62 nM, respectively), rmAb23 neutralized only G614 and Delta (IC_50_: 4.44 and 14.45 nM) while XG014 efficiently neutralized G614, Alpha, Beta and Delta (IC_50_ values from 0.24 to 2.34 nM) but poorly neutralized Omicron. DXP-604 neutralized G614 (IC_50_: 0.3 nM) and all VOCs tested (IC_50_ values from 2.63 to 22.93 nM) although it was less effective against Omicron BA.1 and BA.5 (**Fig. 2C**).

The binding and neutralization assay results are in accordance with structural models showing that certain mutations in the RBDs of VOCs are located in the binding epitope of 01A05 and rmAb23 (**fig. S3, A and B**). However, mutations in RBD residues within the binding epitope of DXP-604 resulted in a less pronounced decrease in neutralization activity including against Omicron BA.1, BA.2 and BA.4/5 which harbour more than 6 RBD substitutions within the mAb epitope (**fig. S3C)**. The epitope recognized by XG014 is mostly outside the hotspots of the RBM, where prevalent mutations are located, explaining the high and broad neutralization potency against Alpha, Beta, and Delta (**fig. S3D**) *(37)*. The apparent resistance of DXP-604 to SARS-CoV-2 mutations was confirmed in a S-pseudotype vesicular stomatitis virus (VSV) neutralization assay showing its potent neutralizing effect against 15 known SARS-CoV-2 variants (IC_50_ from 0.03 nM to 1.6 nM) and other clade 1b sarbecoviruses circulating among other species, including RaTG13 (IC_50_: 0.01 nM) and Pangolin-GD *(38)* (IC_50_: 0.05 nM) (**fig. S4**). Therefore, among the four antibodies analyzed, DXP-604 appeared to be the most broadly effective neutralizer, even when a few residues in its binding epitope had been substituted.

Further analysis of the X-ray crystal structure *(33)* revealed that the footprint of the DXP-604 heavy chain on the RBD is similar to that of LY-CoV016 (etesevimab) but a higher degree of overlap exists between that of the DXP-604 light chain and ACE2-contact surface (**Fig. 2D**). Hydrogen bonds were also formed between S30/S67 in the light chain of DXP-604 and RBD Q498, which is a key ACE2-binding site *(39)*, and between the main chain groups of Q27/S28 and RBD G502 (**Fig. 2E**) *(2)*. In addition, based on the structural model, one single DXP-604 Fab bound to the S-trimer with the 3-RBD in the up position can prevent the binding of ACE2 to all three S monomers (**fig. S2C**). Thus, in contrast to 01A05, rmAb23 and other class I antibodies, the high binding affinity and ACE2-mimicking epitope enable DXP-604 to exhibit a higher tolerance for RBD substitutions and to retain a broad neutralization activity.

### Conversion of monoclonal IgG to IgA1 antibodies increases the neutralization potency

As sIgA1 is the main immunoglobulin type in the upper respiratory tract, the four monoclonal IgG antibodies were engineered as monomeric (mIgA1), dimeric (dIgA1, via coexpression of the J chain), and secretory (sIgA1, by coexpression of the J chain and SC) IgA1 antibodies to compare the binding and neutralizing properties of the various forms of IgA1 (**Fig. 3, A and B**).

**Fig. 3.**
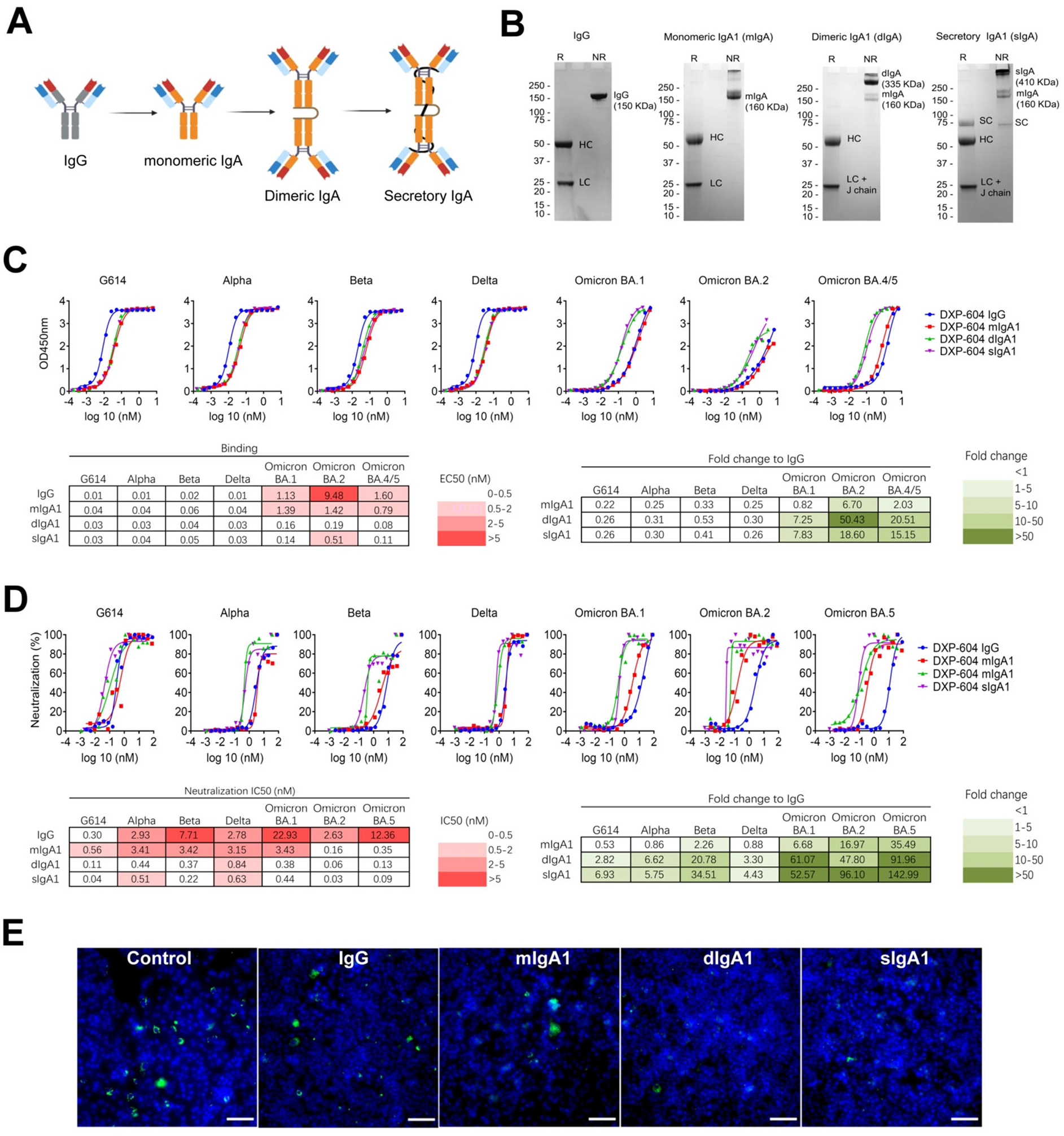
Dimeric and secretory IgA1 showed enhanced binding and neutralization activity against variants of concern (VOCs). (**A**) Illustration showing antibodies engineered from IgG into monomeric, dimeric and secretory IgA1. (**B**) SDS–PAGE under reducing (R) and nonreducing (NR) conditions showing the assembly and purity of DXP-604 IgG and IgA1 antibodies. HC, heavy chain; LC, light chain; SC, secretory component; J chain, joining chain. The J chain migrates at the same molecular weight as the light chain. (**C** and **D)** Binding to RBD (C) and neutralization (D) of G614 and variants of concern (VOCs) by DXP-604 IgG and IgA (monomeric (mIgA1), dimeric (dIgA1) and secretory IgA1 (sIgA1)) antibodies. The EC_50_ and IC_50_ and fold-change differences between IgG and IgA1 antibody forms are indicated. (**E)** Staining of virus following infection of Vero E6 cells with SARS-CoV-2 Omicron BA.1 preincubated with 3.3 nM DXP-604 IgG or IgA1 forms. Omicron BA.1-infected cells were used as a negative control (control). SARS-CoV-2 virus was visualized using Alexa 488 (green)-conjugated antibody, and the nucleus was stained with DAPI (blue). Scale bar, 100 μM. See also fig. S5 and S6.

The conversion of IgG to IgA1 did not strongly increase the binding affinity of the antibodies for the RBD, with generally less than twofold increase as measured by ELISA (**Fig. 3D and fig. S6**). However, dimerization of IgA1 greatly increased the binding of DXP-604 antibodies to the RBD of Omicron BA.1, BA.2 and BA.4/5 (EC_50_ from 0.08 to 0.51 nM) from 7.3- to 50.4- fold compared to those of the parental IgG antibodies (**Fig. 3C**). In addition, XG014 dimeric and secretory IgA1 bound to the BA.2 and BA.4/5 RBDs (EC_50_ from 0.07 to 0.13 nM) 4.7- to 9.9-fold more efficiently than the parental IgG antibodies **(fig. S5C)**.

Switching from IgG to IgA1 and dimerization further increased antibody neutralization activity against real virus, and the effect was greatest for DXP-604, 01A05, and rmAb23 (**Fig. 3D and fig. S6**). DXP-604 monomeric IgA1 showed increased neutralization activity against BA.1 (IC_50_: 3.43 nM, by 6.7-fold), BA.2 (IC_50_: 0.16 nM, by 17.0-fold) and BA.5 (IC_50_: 0.35 nM, by 35.5-fold) (**Fig. 3D**). More importantly, DXP-604 dimeric and secretory IgA1 showed increased neutralization activity against all variants but particularly against the Omicron lineages BA.1, BA.2 and BA.5, which was 47.8- to 143-fold higher than the parental IgG and 2.7- to 9.0-fold higher than the monomeric IgA1 (**Fig. 3, D and E**). DXP-604 dimeric and secretory IgA1 neutralized Omicron BA.1 (IC_50_: 0.38 and 0.44 nM, respectively), BA.2 (IC_50_: 0.06 and 0.03 nM, respectively) and BA.5 (IC_50_: 0.13 and 0.09 nM, respectively) to a potent level that is similar to the neutralization of G614 (IC_50_: 0.11 and 0.04 nM, respectively) and to the counterpart IgG antibodies against G614 (IC_50_: 0.30 nM) (**Fig. 3D**).

An increase in neutralization activity by mIgA1 compared to parental IgG was also observed for 01A05 against G614 (IC_50_: 0.04 nM, by 4.4-fold) and Alpha (IC_50_: 0.93 nM, by 11.2-fold), and for rmAb23 against G614 (IC_50_: 0.89 nM, by 5.0-fold), Alpha (1.66 nM, by >100.6-fold) and Delta (IC_50_: 3.42 nM, by 4.2-fold) (**fig. S6, A and B)**. Dimerization also improved the neutralizing activity of 01A05 and rmAb23 against G614, Alpha and Delta by 4.5- to >179-fold compared to IgG and up to 12.0-fold compared to mIgA1 **(fig. S6, A and B)**. Interestingly, switching to IgA1 and dimerization rescued the neutralizing activity of rmAb23 against Alpha, to some extent (IC_50_: 1.79 and 0.93 nM, for dIgA1 and sIgA1, respectively) and XG014 against Omicron BA.1 (IC_50_: 3.29 and 4.44 nM, for dIgA1 and sIgA1, respectively) (**fig. S6, B and C**). These results suggest that the conversion of IgG to IgA1, particularly to the dimeric and secretory forms of IgA1, significantly improved the neutralizing potency of the antibodies against various VOCs.

### Increased neutralizing potency of dimeric IgA1 is associated with increased avidity

Enhanced neutralization of dimeric IgA1 antibodies exhibited different patterns, suggesting that the epitope and affinity, in addition to valency, may affect their potency *(40)*. The increase in neutralization potency was more profound against variants for which the parental IgG showed lower (but presence of) neutralizing activity. In fact, the fold-change improvement in binding and neutralizing activity of the IgA1 forms was positively correlated with the EC_50_ values of the parental IgG (**fig. S7A to D**) *(28)*. For example, DXP-604, which showed a low RBD binding (EC_50_: 1.60 nM) and a low neutralizing activity (IC_50_: 12.36 nM) against BA.5 as an IgG, was found to be 20.5-fold more potent in binding the RBD and 92.0-fold more potent in neutralizing BA.5 as a dIgA1 (**fig. S7, B and D**). In addition, for DXP-604, the fold-change increase in RBD binding correlated with the fold-change increase in neutralization activity, suggesting that an increase in neutralizing activity is associated with an increased affinity for the RBD, at least for this antibody for which correlation analysis could be individually performed due to broad neutralization (**fig. S7, E and F**).

As the EC_50_ value obtained by ELISA does not measure affinity, and since DXP-604 dimeric and secretory IgA1 neutralized Omicron including the circulating BA.5 lineage with a high potency, we further characterized the binding properties of this antibody. The apparent increase in avidity of antibodies could be due to inter- or intra-S cross-linking. However, according to *in silico* models, only one DXP-604 Fab can bind to each S due to steric hindrance caused by the light chain, thus preventing intra-S binding (**fig. S2C**). We used an avidity assay by surface plasmon resonance (SPR) to experimentally confirm that the dimeric DXP-604 antibody can simultaneously engage two RBDs on different S proteins *(41)*. The association rate, measured as a control, was not affected by avidity and, indeed, was not affected by antigen concentration (**Fig. 4A**). A slower dissociation rate was observed for dimeric IgA1 DXP-604 at higher concentrations of immobilized antigen, consistent with intermolecular avidity effects (**Fig. 4, B and C**). At lower concentrations of immobilized antigen, the *k*_d_ of dIgA1 was faster as intermolecular binding was prevented by the increased distance between RBD molecules, which resulted in loss of avidity (**Fig. 4D**). The S-trimers on the surface of SARS-CoV-2 float readily and are widely spaced (a mean of 24 trimeric S protein per virus) *(42)* at an average distance of 25 nm *(43)*. Theoretically, IgG and mIgA1 antibodies may bind the RBD on two different S-trimers spaced by ∼2 nm. Structural simulations show that dIgA1 and sIgA1 can bridge upon S-trimers ∼11 nm apart (**Fig. 4E**). It is plausible that the dimeric forms are more likely to engage in intermolecular binding on the viral surface (**Fig. 4E**). Thus, the increased neutralization potency of dIgA1 and sIgA1 appears to be, at least partly, due to increased avidity mediated by inter-S-trimer binding on the viral surface but other mechanisms such as aggregation, may also be involved.

**Fig. 4.**
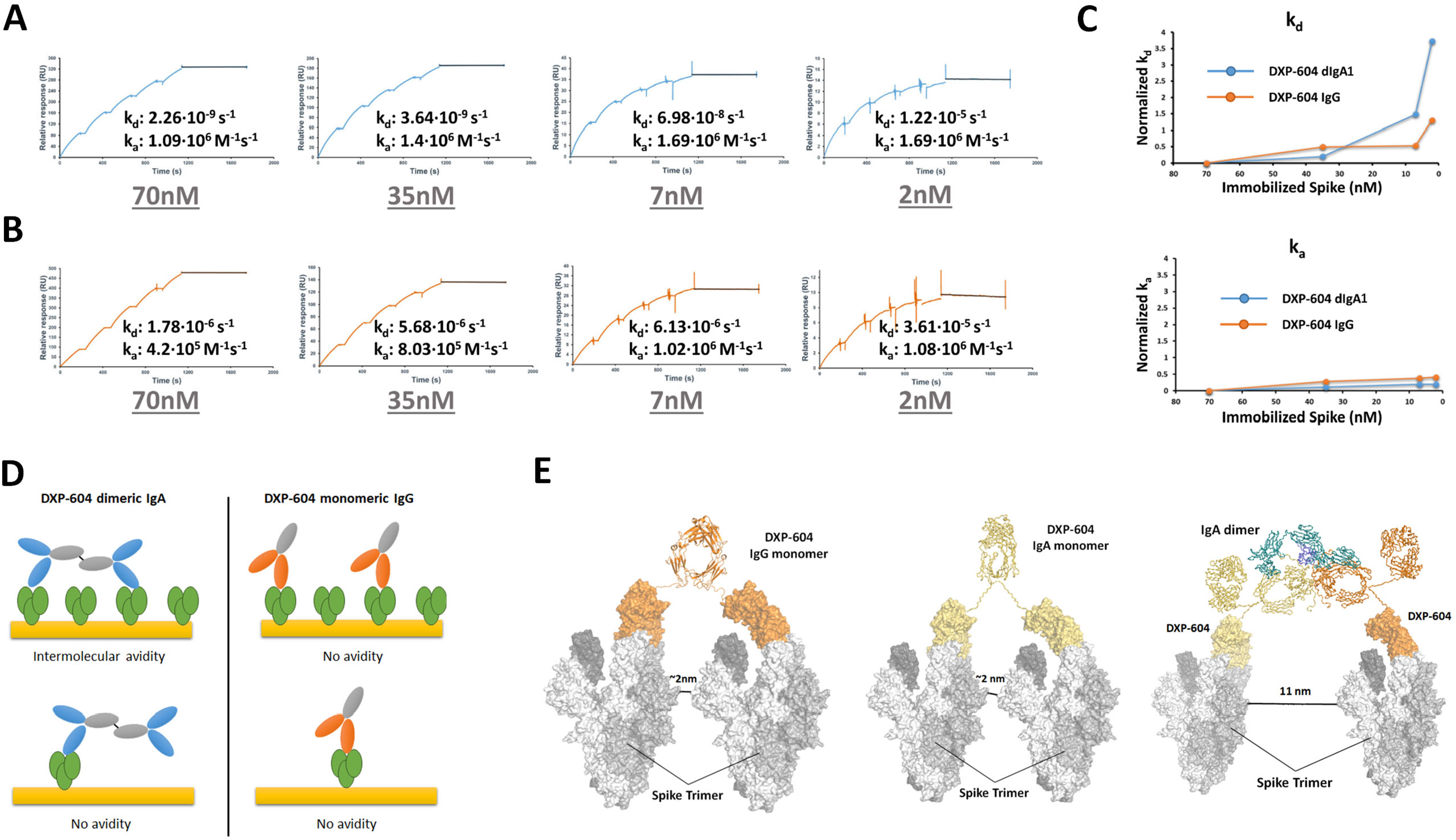
Increased neutralization potency of DXP-604 dimeric IgA is associated with increased avidity. (**A** and **B**) Representative SPR traces used to determine *k*_a_ and *k*_d_ of dimeric IgA1 (A) and monomeric IgG (B) at different concentrations (70, 35, 7 or 2 nM) of immobilized S-trimers on the SPR chip. The *k*a and *k*d are indicated below each graph. (**C**) Plots of normalized *k*a (top) and *k*d (bottom) obtained with different concentrations of immobilized S-trimers. Increasing normalized *k*d values indicate loss of avidity. *k*a and *k*d were normalized against the values at the highest S-trimer concentration. (**D**) DXP-604 dimeric IgA1 and monomeric IgG have different binding modes that are available when high or low quantities of S-trimers are immobilized on the surface of the SPR chip. (**E**) Computational simulation for inter-S linking by DXP-604 monomeric IgG and IgA1, and dimeric IgA1 antibodies. The predicted distance between S-trimers necessary for linking is indicated. See also fig. S7.

### Dimeric IgA is protective against Omicron BA.5 in a mouse model

We subsequently evaluated the protective efficacy of DXP-604 dIgA in transgenic mice expressing human ACE2 (hACE2) infected with Omicron BA.5 using therapeutic and prophylactic intranasal administration (**Fig. 5A**). The viral load in the lungs and tracheal tissues of untreated mice (only PBS) was 10^6.37^ copies and 10^4.29^ copies/g, respectively, at 3 days post infection (dpi). A single intranasal dose of DXP-604 dIgA (60 µg, ∼2.2 mg/kg), 2 hours after challenge with Omicron BA.5, significantly reduced the mean viral load to 10^3.22^ copies/g (a 3.15 log10-fold decrease) in lung tissues and to undetectable levels (a 4.29 log_10_-fold decrease) in tracheal tissues of all three treated mice at 3 dpi (**Fig. 5B**). For prophylactic treatment, administration of 60 µg of DXP-604 dIgA (∼2.9 mg/kg) 4 hours before challenge with the virus, significantly reduced the mean viral load to 10^3.66^ copies/g (a 2.71 log_10_-fold decrease) in lung tissues and to undetectable levels (4.29 log_10_-fold decrease) in tracheal tissues of all treated mice (n = 3) at 3 dpi (**Fig. 5B**). A lower prophylactic dose (40 µg, ∼1.5 mg/kg) significantly reduced the viral load in the trachea only (a 0.84 log_10_-fold decrease). No significant difference in weight change was observed between the treatment and control groups during the experiment (**Fig. 5C**). Together, our data show that prophylactic and therapeutic intranasal administration of DXP-604 dIgA is highly protective against Omicron infection in the respiratory tract.

**Figure 5.**
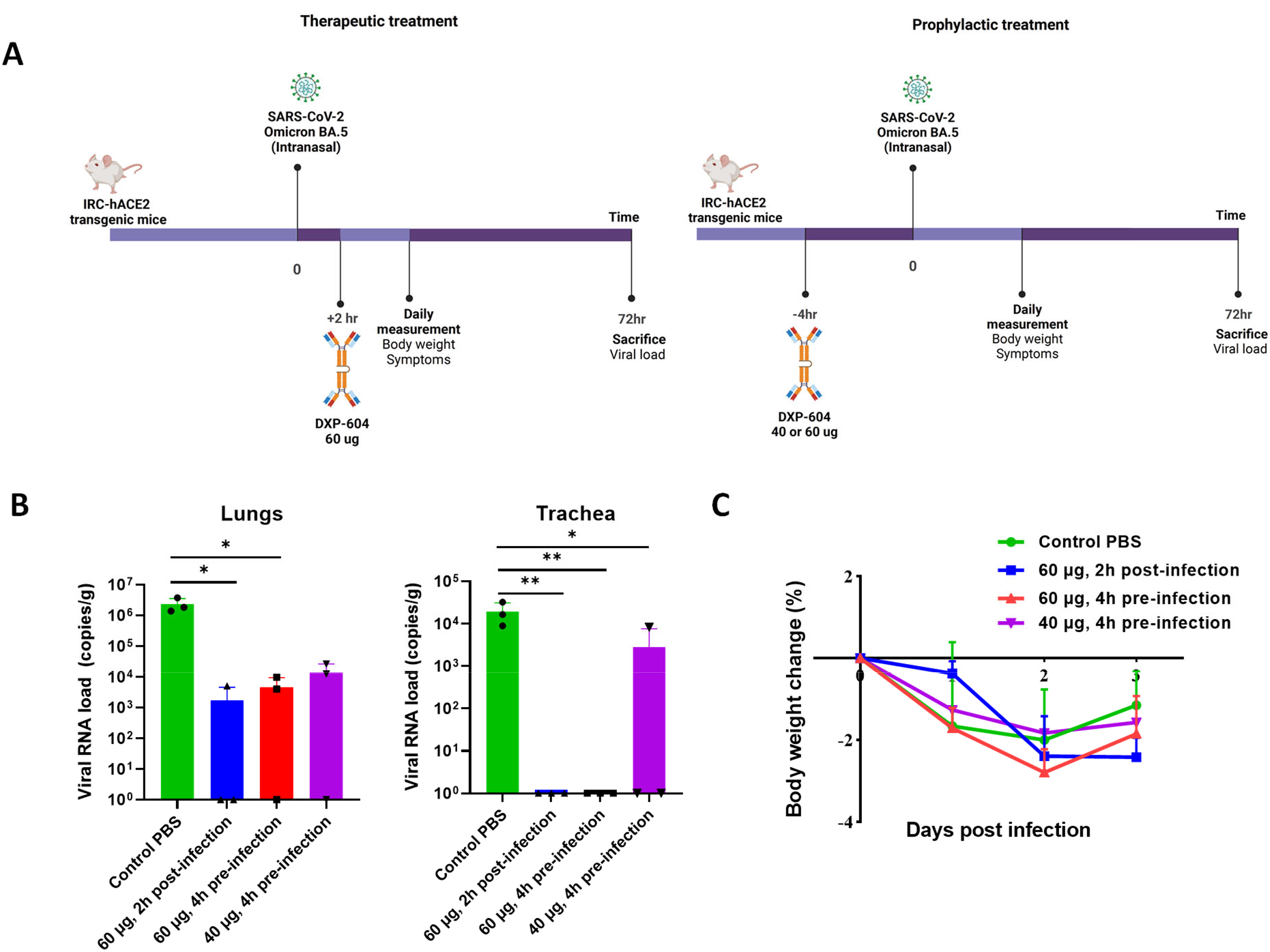
Intranasal administration of dimeric IgA in hACE2 mice is protective against Omicron BA.5. (**A**) Experimental design for the evaluation of DXP-604 dIgA using therapeutic and prophylactic intranasal administration. (**B** and **C**) Viral loads in the lung and tracheal tissues (B), and body weigh change (C) of Omicron BA.5-infected mice after administration of a single dose of DXP-604 dIgA in a therapeutic (60 µg, 2 h post-infection) and prophylactic (40 or 60 µg, 4 h pre-infection) setting. Viral loads and weigh change are expressed as the mean ± standard deviation for three mice. A two-sided unpaired t-test was used. * p < 0.05, and ** p < 0.01.

### Conversion to IgA1 and dimerization increases the neutralization potency against emerging Omicron subvariants

According to the WHO, more than 300 sublineages of Omicron, mainly descendants from BA.2 and BA.5, are circulating globally (https://github.com/gerstung-lab/SARS-CoV-2-International). The BQ.1 and BQ.1.1 subvariants are now dominant in parts of Europe and North America, while BF.7 and XBB are dominant in some regions of Asia but with XBB.1.5 becoming dominant in the United States. Compared to BA.5, BQ.1 carries two additional mutations (K444T and N460K) in the RBD, while BQ.1.1 carries an additional RBD mutation (R346T). BA.2.75 carries three mutations (G446S, N460K, and rev R493Q) in the RBD compared to BA.2 and BA.2.75.2 contains two additional mutations (R346T and F486S). The S protein of XBB has 14 mutations in addition to those found in BA.2, including 9 in the RBD (G339H, R346T, L368I, V445P, G446S, N460K, F486S, F490S and reverse (rev) R493Q), whereas XBB.1 has an additional G252V mutation and XBB.1.5 has the mutation S486P.

Since authentic viruses of the new subvariants were not yet available in our laboratories at the time of the study, we evaluated the neutralization activity of DXP-604 IgG and IgA1 against circulating subvariants, in addition to BA.1, BA.2 and BA.4/5, using pseudovirus assays. Dimeric and secretory IgA1 (IC_50_: < 0.0006 to 0.011 nM) improved the neutralization of BA.1, BA.2 and BA.4/5 Omicron subvariant pseudoviruses by 70.9- to 220-fold compared to monomeric IgG (IC_50_: 0.091 to 0.836 nM) and by 2.0 to 11.5-fold compared to monomeric IgA1 (IC_50_: 0.007 to 0.022 nM) (**fig. S8**). Furthermore, DXP-604 dIgA1 and sIgA1 increased the neutralizing activity against BQ.1 (IC_50_: 3.20 and 1.41 nM, by 2.6- and 6.0-fold), BQ.1.1 (IC_50_: 4.88 and 1.59 nM, by 1.8- to 5.5-fold) and BA.2.75 (IC_50_: 0.028 and 0.013 nM, by 37.0- and 78.0-fold) compared to IgG (**fig. S8**). However, DXP-604 IgA1 forms could not restore neutralization activity against BA.2.75.2 and XBB.1.

Interestingly, 01A05 IgG, which showed no binding to BA.2 (**fig. S5A)**, neutralized BA.2.75.2 (IC_50_: 12.08 nM) and the effect was greatly improved using mIgA1 (IC_50_: 0.067 nM, by 180- fold), dIgA1 (IC_50_: 0.0165 nM, by 733-fold) or sIgA1 (IC_50:_ 0.006 nM, by 2023-fold) (**Fig. S9**). Furthermore, an increase in neutralization activity (by >200-fold) against XBB.1 was observed for 01A05 mIgA1 (IC_50_: 0.30 nM), dIgA1 (IC_50_: 0.13 nM) or sIgA1 (IC_50:_ 0.18 nM) compared to IgG (IC_50:_ >66 nM), which poorly neutralizes this variant. As previously observed with a few other monoclonal antibodies *(44, 45)*, the R493Q reversion mutation in BA.2.75.2 and XBB.1 at least partially restored the 01A05 neutralizing epitope found in the ancestral SARS-CoV-2 (**fig. S3A**).

Since many Omicron subvariants harbouring multiple convergent mutations are simultaneously circulating in different parts of the world *(12)*, our results suggest that a cocktail of dimeric or secretory IgA1 antibodies, including DXP-604 or 01A05 described here and those newly identified broad neutralizers *(12, 46, 47)*, would be necessary to neutralize most, if not all, emerging Omicron subvariants and future VOCs.

## DISCUSSION

SARS-CoV-2 primarily infects the upper respiratory tract, where the mucosal immune response is expected to be mainly induced in the nasopharynx, via the tonsils and adenoids, collectively referred to as the nasopharynx-associated lymphoid tissue (NALT). Systemic immunization has generally been considered ineffective in generating protective mucosal immune responses, although certain antigen and adjuvant combinations can elicit mucosal immune responses, including sIgA antibodies *(48)*. The mechanism of this induction remains poorly understood and may involve the migration of vaccine antigens, antigen-loaded antigen-presenting cells and/or antigen-specific B cells to mucosal-associated lymphoid tissues (MALT) *(48–50)*. In this study, the inactivated and mRNA vaccines induced salivary IgA anti-RBD antibodies in 60% of individuals but at a low level in comparison to patients with BTI, particularly against Omicron. Furthermore, additional vaccine doses did not boost the salivary RBD-specific IgA response and the antibody levels gradually decreased over time following the third dose of mRNA vaccine, whereas BTI gave rise to a higher IgA response with no decrease over the time monitored thus far (up to 9 months). We previously showed that the level of mucosal RBD-specific IgA antibodies induced by vaccination might be associated with protection against subsequent BTI *(31)*. These findings support the importance of the mucosal immune system in the defence against SARS-CoV-2 and the need for the development of vaccines that can elicit a stronger mucosal IgA response *(20, 27, 51)* and a therapy based on intranasally administered antibodies.

Omicron lineages and subvariants are currently circulating worldwide, and they carry mutations that confer resistance to most potent neutralizing antibodies, including those clinically approved *(12, 52)*. In agreement with previous studies, we show that only DXP-604, one of our identified SARS-CoV-2 RBD-neutralizing antibodies encoded by IGHV3-53, can broadly neutralize most SARS-CoV-2 variants and selected subvariants *(12)*. This outcome is probably achieved through high affinity and ACE2-mimicking interactions via light chain complementarity-determining regions (CDRs), especially hydrogen bonds formed with RBD G502 and Q498. Although the neutralizing activity of DXP-604 IgG against Omicron BA.1, BA.2 and BA.5 *(2, 52)* was lower than that of LY-CoV1404 (bebtelovimab) *(52, 53)* and other IgG antibodies in preclinical or clinical development *(54–58)*, DXP-604 dimeric and secretory IgA1 showed a 50- to 150-fold increase in potency against the BA.1, BA.2 and BA.5 variants (IC_50_: 0.03 to 0.44 nM), reaching the level of one of the most potent antibodies, LY-CoV1404 (IC_50_ 0.08-0.1 nM) (**table S2**). Contrary to LY-CoV1404 and other antibodies approved for clinical use *(5, 12)*, DXP-604 dimeric and secretory IgA1 could also neutralize BQ.1 and BQ.1.1, which make up the largest share of new COVID-19 infections in Europe and North America. However, DXP-604 was recently shown not to be able to neutralize sublineages carrying the mutation F486S, explaining its inability to neutralize BA.2.75.2 and XBB.1 *(12)*. Nevertheless, 01A05 dimeric and secretory IgA1 strongly improved neutralization potency against BA.2.75.2 and XBB.1, suggesting that a combination of dimeric or secretory IgA1 antibodies could be used to neutralize a wider range of variants with higher potency.

The subclass proportions vary with mucosal site but typically range from 80 to 90% IgA1 in nasal and male genital secretions, 60% IgA1 in saliva, to 60% IgA2 in colonic and female genital secretions *(23)*. IgA1 and IgA2 differ mostly in the hinge region, which is significantly longer (13 aa) in IgA1. As a result, IgA1 is more like a T-shaped molecule *(59)*, whereas IgA2 resembles the more rigid classical Y-shape of IgG *(60)*, as revealed by X-ray crystal structures. Consistent with previous studies, a switch to monomeric IgA may result in an increase in the neutralizing capacity of certain antibodies, such as DXP-604, and against some variants (particularly BA.1, BA.2 and BA.5) *(61)* compared to that of IgG. The longer IgA1 hinge may confer increased flexibility and allow two Fabs to reach two RBDs in the trimer at the same time, thus enhancing SARS-CoV-2 neutralization *in vitro* compared to that by their IgG counterparts. However, computational modelling showed that although 01A05 and XG014 Fabs each simultaneously can bind two RBDs on the same trimeric S protein, DXP-604 Fabs cannot bind more than one site due to steric hindrance caused by the light chain (**fig. S2**). Alternatively, the increased flexibility in the hinge region could help the binding of one Fab fragment to RBD or promote linking of S on two different virus particles. We tested DXP-604 antibodies in the IgA2 format and found that DXP-604 mIgA1 neutralizes Omicron BA.2 and BA.5 with more than 15-fold higher potency than DXP-604 mIgA2 (unpublished data). However, considering the higher proportion of IgA2 in the intestinal tract and its increased resistance to bacterial proteases due to shorter hinge, dimeric or secretory IgA2 might still be useful for oral delivery to prevent faecal-oral transmission or reduce gastrointestinal symptoms *(62)*.

Intranasal delivery of IgM (3.5 mg/kg) *(40)* and IgG (3 mg/kg) *(63)* antibodies after infection with SARS-CoV-2 (Beta or Omicron BA.2) was previously shown to reduce the viral load in the lungs of hACE2 expressing mice. We showed here that, for the first time, a therapeutic (2.2 mg/kg) or prophylactic (2.9 mg/kg) intranasal administration of dIgA, confer significant protection against Omicron BA.5 in the hACE2 transgenic mouse model and considerably reduced the viral load in both the lung and trachea. The higher conferred protection in the trachea compared to the lung could be due to the biodistribution of the antibodies following intranasal administration. Thus, intranasal delivery of dimeric or secretory DXP-604 IgA in combination with other IgA antibodies directly to the site of infection may be an effective approach to achieve immediate protection against SARS-CoV-2 infection, which is needed at a small window for intervention, such as prevention in high-risk settings or postexposure prophylaxis. IgA does not activate complement and may inhibit complement activation induced by IgG and IgM, thus reducing inflammation *(19)*.

Through an increase in avidity and possibly aggregation, the dimerization of IgA can considerably increase the potency of broadly neutralizing antibodies, including against emerging variants with multiple RBD escape mutations such as Omicron, and offer effective protection of the respiratory tract in animal models. A formulation of dIgA or sIgA anti-SARS-CoV-2 antibodies may eventually be applied by susceptible individuals themselves with or without medical supervision, and thus has a wider coverage, which is especially important in resource-poor areas. Nasal delivery of antibodies is likely to be carried out using low doses, since the drug is administered locally, where it is most needed, in contrast to the larger amounts needed for systemic application. To ensure sustainable production of the therapeutic agent, these antibodies can be expressed in plants (rice, tobacco) *(64)*. Produced in this way, recombinant sIgA-based passive immunotherapies or prophylactics could represent extremely effective tools for the control of SARS-CoV-2 infections.

## MATERIALS AND METHODS

### Study Design

The objectives of the study were to evaluate the mucosal immune response to SARS-CoV-2 in vaccinated individuals and to develop dimeric and secretory IgA1 antibodies for mucosal prophylactic and therapeutic treatment. We initially tested the presence and dynamics of salivary antibodies against the RBDs of G614 (wild-type) and Omicron lineages BA.1, BA.2, and BA.4/5 in saliva and plasma of individuals with different types of vaccinations and followed the dynamics of salivary IgA and IgG antibody responses over time (up to 9 months). The anti-RBD antibody levels and total IgA immunoglobulin levels were measured by ELISA.

We subsequently engineered and characterized recombinant monomeric, dimeric and secretory IgA1 antibodies derived from four neutralizing IgG monoclonal antibodies (mAbs) (1A05, rmAb23, DXP-604 and XG014) targeting the RBD domain of the spike protein. The anti-RBD IgG and IgA1 antibody forms were produced and tested for binding to RBD in ELISA and for neutralizing activity using authentic virus (G614, Alpha, Beta, Delta, Omicron BA.1, BA.1, BA.5) and pseudovirus (Omicron BA.1, BA.2, BA.5, BQ.1, BQ.1.1, BA.2.75, BA.2.75.2, and XBB.1). Computational modelling for predicting the antibody structure and antibody-RBD interaction was performed. We used an avidity assay by surface plasmon resonance (SPR) to experimentally confirm that the dimeric IgA antibody can simultaneously engage two RBDs on different S proteins. We also evaluated the protective efficacy of DXP-604 dimeric IgA in a mouse model using intranasal therapeutic and prophylactic administration.

### Study subjects

Study inclusion criteria included subjects who were older than 18 years of age, received inactivated and/or mRNA vaccines with a documented vaccination history (type of vaccine, number of doses, interval between the doses, days after the latest dose, and infection data), and were willing and able to provide written informed consent. The study included 185 samples from 111 healthy volunteers (66% females, median age of 30 years) in Sweden in 2021-2022 who received two or three doses of inactivated vaccine (CoronaVac, Sinovac or BBIBP-CorV, Sinopharm), 1 to 3 doses of an mRNA vaccine (BNT162b2, Pfizer–BioNTech or mRNA-1273, Moderna) or a combination of both (two doses inactivated vaccine followed by a heterologous mRNA boost), some of whom had experienced BTIs during the Omicron BA.1 wave. A group receiving one or two doses of mRNA vaccine after SARS-CoV-2 infection (during the G614 wave) was also included (**table S1**). Samples were collected 5-59 days (median day 20) after each mRNA dose including after mRNA heterologous boost, 6-92 days (median day 48) after doses 2 and 3 of inactivated vaccine and 8-43 days (median day 17.5) after BTI (**table S1**). For a subset of individuals, we also followed the dynamics of the antibody response 9 months after the second and third doses of mRNA vaccinations and 10 months after BTI in individuals who received 2 or 3 mRNA vaccine doses (**table S1**). Infection was confirmed when an individual tested positive for antigen or qPCR test. Saliva and plasma samples from prevaccinated, uninfected healthy donors in our cohort were also collected and used as negative controls (n=7). The study was approved by the ethics committee of the institutional review board of Stockholm.

### Detection of antibodies specific to SARS-CoV-2 RBD

To assess anti-RBD binding activity, high-binding Corning half-area plates (Corning #3690) were coated with RBD derived from G614, BA.1, BA.2 or BA.4/5 (1.7 μg/ml) in PBS and incubated overnight at 4°C. Serial dilutions of saliva in PBS with 5% skim milk supplemented with 0.1% Tween 20 were added, and the plates were subsequently incubated for 1.5 hours at room temperature. The plates were then washed and incubated for 1 hour with horseradish peroxidase (HRP)-conjugated goat anti-human IgM (Invitrogen #A18835), goat anti-human IgA (Jackson #109-036-011), or goat anti-human IgG (Invitrogen #A18805) antibodies (all diluted 1:5000 in PBS supplemented with 5% skim milk and 0.1% Tween 20). For detection of secretory immunoglobulins (sIgA and sIgM), plates were incubated for 1 hour with HRP-conjugated goat anti-secretory component antibodies (Nordic–MUbio, #GAHu/SC/PO). The bound antibodies were visualized using tetramethylbenzidine substrate (Sigma #T0440). The colour reaction was stopped with 0.5 M H_2_SO_4_ after 10 min of incubation, and the absorbance was measured at 450 nm in an ELISA plate reader (Varioskan, Thermo Scientific).

Plasma IgA and IgM and salivary antibody levels are reported as arbitrary units (AU)/ml based on a standard curve generated with data derived from a serially diluted highly positive in-house serum pool. Plasma and salivary IgG levels are expressed as binding antibody units (BAU)/ml after calibrating in-house standards to the WHO International Standard for anti-SARS-CoV-2 immunoglobulin (NIBSC, 20/136) *(65, 66)*. For secretory immunoglobulin, serial dilutions of human monoclonal secretory IgA anti-RBD antibodies (DXP-604) were used for the generation of a standard curve and measurement of concentrations (ng/ml). Salivary IgA anti-RBD antibodies were normalized according to the total level of salivary IgA (AU/µg total IgA) to compensate for the different salivary flow rates between individuals. The positive cut-off was calculated to be 2 standard deviations (2SD) higher than the mean of a pool of samples taken from prevaccinated and noninfected individuals (n = 7).

### Detection of total IgA immunoglobulin

To assess total IgA, high-binding Corning half-area plates (Corning #3690) were coated overnight at 4°C with polyclonal goat anti-human IgA (Southern Biotech, #C5213-R466) (2 μg/ml) in PBS. Serial dilutions of saliva in PBS supplemented with 5% skim milk and 0.1% Tween 20 were added, and the plates were subsequently incubated for 1.5 hours at room temperature. The plates were then washed and incubated with HRP-conjugated polyclonal goat anti-human IgA (Jackson #109-036-011) diluted 1:15000. The bound antibodies were detected as described above. Serial dilutions of human monoclonal IgA were used for the generation of standard curves and measurement of concentrations (ng/ml).

### Production of SARS-CoV-2 RBD protein

The RBDs of G614, Alpha, Beta, and Omicron (BA.1, BA.2, BA.4/5) variants were ordered as GeneString from GeneArt (Thermo Fisher Scientific). All sequences of the RBD (aa 319-541 in GenBank: MN908947) were inserted into an NcoI/NotI compatible variant of an OpiE2 expression vector carrying the N-terminal signal peptide of the mouse Ig heavy chain and a C-terminal 6xHis-tag. RBD of G614, Beta, Delta and Omicron were expressed in a baculovirus-free expression system in High Five insect cells and purified on HisTrap Excel columns (Cytiva) followed by size-exclusion chromatography on 16/600 Superdex 200-pg columns (Cytiva) *(67, 68)*.

### Isolation of IgG antibodies by sorting RBD-binding memory B cells

The 01A05 IgG antibody was isolated by sorting RBD-binding memory B cells from convalescent patients infected with the G614 strain. The G614 RBD was labelled with either allophycocyanin (APC) or phycoerythrin (PE) for use in a two-fluorescent-dye sorting strategy.

XG014 *(35)* and DXP-604 *(2, 33)* IgG were previously isolated by sorting RBD-binding memory B cells from convalescent patients infected with the Wuhan strain. rmAb23 was previously isolated using an antibody repertoire prepared by sequencing PBMCs from patients infected with the Wuhan strain followed by matching of the VH3-53-J6 heavy chain with a common IGKV1-9 light chain to produce recombinant antibodies *(32)*.

### Cloning of neutralizing IgG antibodies to generate the IgA forms

The heavy and light chain variable genes of DXP-604, XG014, 01A05 and rmAb23 neutralizing IgG antibodies were cloned into separate pcDNA 3.4 vectors to mediate fusion to an IgA1 constant region and a light chain constant region gene (kappa for 01A05, rmAb23 and DXP-604 and lambda for XG014), respectively (GenScript). The J-chain and SC genes were cloned into separate pcDNA 3.4 expression plasmids for the assembly of dimeric IgA and secretory IgA.

### Production and purification of antibodies

The IgG and IgA1 antibodies 01A05, rmAb23, XG014 and DXP-604 were produced by transfection of HD CHO-S cells with plasmids in a 30-ml volume (GenScript). Monoclonal IgA1 antibodies were produced in CHO cells transiently transfected with two plasmids expressing a heavy and light chain. For the expression of dimeric and secretory IgA1 antibodies, cells were cotransfected with plasmids carrying the J-chain and SC. The IgG and IgA1 antibodies were purified by single-step affinity chromatography using immobilized protein A (MabSelect SuRe™ LX, Cytiva) or anti-IgA antibody (CaptureSelect™ IgA Affinity Matrix), respectively (GenScript).

### Binding of monoclonal and recombinant antibodies specific to SARS-CoV-2 RBD

To assess the anti-RBD IgG or IgA binding activity, high-binding Corning half-area plates (Corning #3690) were coated with RBD derived from G614WT, Beta, Delta and Omicron (BA.1, BA.2, BA.4/5) (1.7 μg/ml) in PBS and incubated overnight at 4°C *(66)*. Serial dilutions of antibody in PBS with 0.1% bovine serum albumin (BSA) were added, and the plates were subsequently incubated for 1.5 hours at room temperature. The plates were then washed and incubated with HRP-conjugated goat anti-human IgG (Invitrogen #A18805) or goat anti-human IgA (Jackson #109-036-011) (diluted 1:15000 in 0.1% BSA-PBS) followed by tetramethylbenzidine substrate. For each sample, the EC_50_ values were calculated using four-parameter nonlinear regression GraphPad Prism 7.04 software *(69)*.

### Pseudovirus neutralization assay based on the VSV platform

The genes encoding the full S protein of D614G, Alpha, Beta, Gamma, Delta, Kappa, Delta plus, Mu, Lambda, Iota, Omicron (BA.1, BA.2, BA.3, BA.1/BA.2 subvariants BA.1.1 (BA.1+R346K), BA.2.12.1 (BA.1+L452Q+S704L) and BA.2.13 (BA.1+L452M)) and clade 1b SARS-CoV-2 related sarbecoviruses (RaTG13 and Pangolin-GD) were cloned into the pcDNA3.1 vector. S pseudotyped virus was prepared based on a VSV pseudotyped virus production system. After transfection and culturing, the supernatant containing pseudotyped virus was harvested, filtered, diluted to obtain the same particle number across samples, as determined based on quantitative analysis by RT‒PCR, and frozen at −80°C for further use. Pseudovirus neutralization assays were performed using the Huh-7 cell line (Japanese Collection of Research Bioresources [JCRB], 0403) or 293T cells overexpressing human angiotensin-converting enzyme 2, also called 293T-hACE2 cells (Sino Biological Company). Monoclonal antibodies were serially diluted in DMEM (HyClone, SH30243.01) and mixed with pseudovirus in 96-well plates. After the mixture was incubated for 1 hour in a 37°C incubator with 5% CO_2_, digested Huh-7 cells or 293T-hACE2 cells were seeded. After incubation, the supernatant was discarded, D-luciferin reagent (PerkinElmer, 6066769) was added to avoid a light reaction, and the luminescence value was detected with a microplate spectrophotometer (PerkinElmer, Ensight, 6005290). The IC_50_ was determined by a four-parameter logistic regression model.

### Pseudovirus neutralization assay based on the HIV platform

The human codon-optimized gene coding for the S protein of G614, BA.1, BA.2 and BA.4/5 lacking the C-terminal 19 codons (S_Δ19_) was synthesized by GenScript. The S_Δ19_ gene of BA.2.75, BA.2.75.2, BQ.1, BQ.1.1 and XBB.1 was constructed by site-directed mutagenesis (QuikChange Multi Site-Directed Mutagenesis Kit, Agilent) using the BA.2 or BA.4/5 S_Δ19_ gene as a template. To generate (HIV-1/NanoLuc2AEGFP)-SARS-CoV-2 particles, three plasmids were used, with a reporter vector (pCCNanoLuc2AEGFP), HIV-1 structural/regulatory proteins (pHIV_NL_GagPol) and SARS-CoV-2 S_Δ19_ carried by separate plasmids as previously described *(70)*. 293FT cells were transfected with 7 µg of pHIV_NL_GagPol, 7 µg of pCCNanoLuc2AEGFP, and 2.5 µg of pSARS-CoV-2-S_Δ19_ carrying the S_Δ19_ gene from G614 or Omicron variants (at a molar plasmid ratio of 1:1:0.45) using 66 µl of 1 mg/ml polyethylenimine (PEI).

Tenfold serially diluted monoclonal antibodies were incubated with pseudovirus carrying the S protein from SARS-CoV-2 (G614, BA.1, BA.2, BA.4/5, BA.2.75, BA.2.75.2, BQ.1, and BQ.1.1) for 1 hour at 37 °C. The mixture was subsequently incubated with 293T-hACE2 cells for analyses of G614 or Omicron pseudoviruses for 48 hours, after which the cells were washed with PBS and lysed with Luciferase Cell Culture Lysis reagent (Promega). NanoLuc luciferase activity in the lysates was measured using the Nano-Glo Luciferase Assay System (Promega) with a Tecan Infinite microplate reader. The relative luminescence units were normalized to those derived from cells infected with pseudotyped virus in the absence of monoclonal antibodies. The IC_50_ values for the monoclonal antibodies were determined using four-parameter nonlinear regression (the least squares regression method without weighting) (GraphPad Prism 7.04 software).

### Microneutralization assay

The SARS-CoV-2 G614 strain and VOCs (Alpha, Beta, Delta, and Omicron BA.1, BA.2 and BA.5) were isolated from patients in Pavia, Italy, and identified by next-generation sequencing. The neutralizing activities of the antibodies were determined via microneutralization assays *(71)*. Briefly, 50 μl of an antibody, starting at 25 µg/ml and increased in a twofold dilution series, was mixed in a flat-bottom tissue culture 96-well microtiter plate (COSTAR, Corning Incorporated) with an equal volume containing a 100 median tissue culture infectious dose (TCID50) of a SARS-CoV-2 strain that had been previously titrated. All dilutions were performed using Eagle’s minimum essential medium to which 1% (w/v) penicillin, streptomycin and glutamine and 5 µg/ml trypsin had been added. After 1 hour of incubation at 33°C in 5% CO_2_, VERO E6 cells (VERO C1008 [Vero 76, clone E6, Vero E6]; ATCC® CRL-1586™) were added to each well. After 3 days of incubation, the cells were stained with Gram’s crystal violet solution (Merck) plus 5% formaldehyde (40% m/v) (Carlo Erba S.p.A.) for 30 min. Microtiter plates were then washed in tap water. Wells were analysed to evaluate the degree of cytopathic effect compared to untreated controls. Each experiment was performed in triplicate. The IC_50_ was determined using four-parameter nonlinear regression (GraphPad Prism).

### Virus was detected by immunofluorescence following neutralizing antibody assay

DXP-604 monomeric IgG and monomeric, dimeric, and secretory IgA1 antibodies, each at a concentration of 3.3 nM, were mixed with an equal volume of Omicron BA.1 (200 plaque-forming units (PFU) per 100 µL) in Dulbecco’s modified Eagle’s medium (DMEM) supplemented with 2% foetal calf serum (FCS) and gentamycin and incubated at 37°C for 1 hour. Next, confluent VeroE6 cells in 96-well plates were washed twice with serum-free DMEM, and the cells were infected with 100 µl of an mAb-virus mix or with virus and no antibodies for 1 hour at 37°C with 5% CO_2_. The cells were washed twice, and 100 µL of DMEM with 2% FCS and gentamycin was added to each well. After 9 hours of infection, the cells were fixed with 4% formaldehyde overnight. The cells were washed once with PBS and permeabilized with 0.2% Triton-X in PBS for 15 min at room temperature. Nonspecific binding was blocked with 3% BSA in PBS at 37°C for 1 hour. A primary antibody (mouse anti-dsRNA) at a 1:200 dilution with PBS containing 2% BSA was added and incubated for 2 hours at 37°C. After 3 washes, secondary goat anti-mouse Alexa 488 (Jackson ImmunoResearch) in 1:200 in PBS containing 1% BSA and DAPI nuclear stain was added and incubated for 1 hour at 37°C. After 4 washes with PBS, 150 µL of PBS was added to each well, and microscopy was performed using a Leica DMi8 with 20X objectives. The same microscopy settings were used for all images. ImageJ version 2.1.0. was used for processing, and all images were linearly stretched equally. The concentration of each antibody analysed (3.3 nM) corresponded to the IC_50_ of the monomeric IgA antibody.

### Surface plasmon resonance (SPR)

Antibody binding properties were analysed at 25 °C using a Biacore 8K instrument (GE Healthcare) with 10 mM HEPES pH 7.4, 150 mM NaCl, 3 mM EDTA, and 0.005% Tween-20 as running buffer. SARS-CoV-2 S-trimers (2, 7, 35 and 70 nM) were immobilized on the surface of a CM5 chip (Cytiva) by standard amine coupling. Increasing concentrations of antibody (3.125, 6.25, 12.5 25, and 50 nM) were injected at a single-cycle kinetics setting (association time: 180 sec, flow rate: 30 µL/min), and dissociation was followed for 10 min. Analyte responses were corrected for nonspecific binding and buffer responses. Curve fitting and data analysis were performed with Biacore^TM^ Insight Evaluation Software v.2.0.15.12933. The fitting of Kon and Koff was performed separately using two different kinetics models. For Kon, a 1:1 binding model was used, while for Koff, a 1:1 dissociation model was used.

### Computational modelling

Computational structure modelling was performed based on the binding of 01A05 to the RBD of variants (Alpha, Beta, Delta, and Omicron) or through previously published information (Protein Data Bank (PDB) files) describing docking (rmAb23) *(32)* or crystallization (DXP-604 and XG014) studies *(33, 34)*.

The 01A05 variable fragment was modelled according to the canonical structure method with the program RosettaAntibody *(72)* as previously described *(73)*. Docking was performed using RosettaDock v3.1 as previously described *(74)*. In summary, the 01A05 model was docked to the WT RBD experimental structure (PDB ID: 6m17). Among the thousands of computationally generated complexes, the decoy in better agreement with experimental data (competition with hACE2 and differential neutralization activity against SARS-CoV-2 variants) was selected and further refined by computational docking.

The selected models of 01A05 and rmAb23 were subjected to a 350 ns molecular dynamics (MD) simulation to adjust the local geometry and verify that the structure was energetically stable. MD was performed with GROMACS *(75)*. The system was initially set up and equilibrated through standard MD protocols: proteins were centred in a triclinic box, 0.2 nm from the edge, filled with SPCE water model and 0.15 m Na^+^Cl^-^ ions using the AMBER99SB-ILDN protein force field. Energy minimization was performed to allow the ions to achieve a stable conformation. Temperature and pressure equilibration steps, respectively at 310 K and 1 Bar, of 100 ps each were completed before performing the full MD simulations with the abovementioned force field. MD trajectory files were analyzed after removal of periodic boundary conditions. The stability of each simulated complex was verified by root mean square deviation and visual analysis.

The structures of mIgG, mIgA and dIgA DXP-604 bound to two SARS-CoV-2 S trimers were built using PyMOL software (The PyMOL Molecular Graphics System, Version 2.0 Schrödinger, LLC).

### Antibody protection in an animal model

The animal study was performed in an animal biosafety level 3 (ABSL3) facility using HEPA-filtered isolators. All animal procedures were approved by the Institute of Laboratory Animal Sciences (ILAS), Chinese Academy of Medical Sciences (CAMS) and Peking Union Medical College (PUMC) (BLL22007).

SARS-CoV-2/human/CHN/GD-5/2022 (Omicron BA.5, GenBank: OP678016) was provided by Guangdong Provincial Center for Disease Prevention and Control. The virus was produced with VERO E6 cells, and the SARS-CoV-2 titer was determined by the TCID_50_ method.

Specific-pathogen-free (SPF) IRC-hACE2 mice, 8-10 weeks old (18-32 g), were provided by the Institute of Medical Experimental Animals, Chinese Academy of Medical Sciences. hACE2 mice were randomly divided into 4 groups with three mice in each group. The animals were inoculated intranasally with 50 µl of authentic SARS-CoV-2 Omicron BA.5 (10^5^ TCID50/ml).

For therapeutic treatment, the mice were administered 60 µg DXP-604 dIgA using a nasal drop (50 µl), 2 hours after viral challenge, and for prophylactic treatments, 40 or 60 µg DXP-604 dIgA was administered 4 hours before challenge. The negative control group received PBS only. The infected mice were observed daily to record body weights and symptoms. The animals were euthanized by exsanguination at 3 days post-infection while under deep anaesthesia. Tissue specimens, including samples from the lung and trachea, were collected for quantification of viral load.

Viral load analysis was performed by qRT-PCR. Lung and trachea homogenates were prepared by using an electric homogenizer. The total RNA of the lungs and trachea was extracted with the RNeasy Mini Kit (Qiagen). Reverse transcription was processed with the PrimerScript RT Reagent Kit (TaKaRa) according to the manufacturers’ instructions. qRT-PCR reactions were performed using the PowerUp SYBG Green Master Mix Kit (Applied Biosystems), according to following cycling protocol: 50°C 2 min, 95°C 2 min, followed by 95°C 15 s, 60°C 30 s for 40 cycles, and then by the melting curve analysis: 95°C 15 s, 60°C 1 min, 95°C 45 s. Forward primer 5′-TCGTTTCGGAAGAGACAGGT-3′ and reverse primer 5′-GCGCAGTAAGGATGGCTAGT-3′ were used in qRT-PCR. Standard curves were constructed by using 10-fold serial dilutions of recombinant plasmids with known copy numbers (from 1.47 × 10^9^ to 1.47 × 10^1^ copies/μl).

### Quantification and statistical analysis

A two-sided Mann‒Whitney U test was performed for comparisons of anti-SARS-CoV-2 antibody levels between groups. A Wilcoxon signed-rank test was used for comparison of paired samples. Correlation analysis between antibody levels was performed using Spearman’s rank correlation. A two-sided t-test was used to compare the viral load in the mouse model. All analyses and data plotting were performed with GraphPad 7.05. A p value less than 0.05 was considered to be statistically significant.

### Supplementary Materials

Figs. S1 to S9

Tables S1 and S2

Data file S1

## Acknowledgements

We thank Prof. Michel Nussenzweig (Rockefeller University), for sharing reagents and insightful discussions.

## Author contributions

H.M., L.H., and Q.P.H. conceived the study. Y.C., F.Z., L.SI., J.C.S., M.P., R.S., I.C., M.HA., A.P., J.Y., L.D., E.P., F.BE., M.S., C.G., N.S., S.B., N.R., M.K.B.,Y.X., C.G., L.K., L.CA., Q.W., M.HU., Z.S., L.F., L.SV., L.CH., M.C.N., F.BA., C.Q., J.X., L.H., X.L.Y. and L.V. were involved in the sample collection, antibody isolation, protein purification, binding assays, surface plasmon resonance, and neutralization assays. L.B and C.Q. tested the antibodies in an animal model. C.G., A.CA. and D.F.R. provided unpublished reagents. H.M., Y.C., F.Z., L.SI., J.C.S., M.P., R.S., I.C., and X.L.Y. were involved in the interpretation of the raw data. F.Z., Y.C., L.SI., M.P., R.S., M.HA., and H.A. performed the computational analysis and prepared the images. H.M., Y.C., L.H., X.L.Y., L.V., X.S.X. and Q.P.H. designed the laboratory protocols and supervised the project. H.M., Y.C., L.H., X.S.X., and Q.P.H. wrote the original draft. All authors revised and approved the paper.

## Data and materials availability

The data that support the findings of this study are available within the Article and Supplemental data. All other data are available from the corresponding author upon reasonable request.

## REFERENCES

1. Y. Chen, Q. Liu, L. Zhou, Y. Zhou, H. Yan, K. Lan, Emerging SARS-CoV-2 variants: Why, how, and what’s next? Cell Insight 1, 100029 (2022).

2. Y. Cao, J. Wang, F. Jian, T. Xiao, W. Song, A. Yisimayi, W. Huang, Q. Li, P. Wang, R. An, J. Wang, Y. Wang, X. Niu, S. Yang, H. Liang, H. Sun, T. Li, Y. Yu, Q. Cui, S. Liu, X. Yang, S. Du, Z. Zhang, X. Hao, F. Shao, R. Jin, X. Wang, J. Xiao, Y. Wang, X. S. Xie, Omicron escapes the majority of existing SARS-CoV-2 neutralizing antibodies. Nature 602, 657–663 (2022).

3. A. Tuekprakhon, R. Nutalai, A. Dijokaite-Guraliuc, D. Zhou, H. M. Ginn, M. Selvaraj, C. Liu, A. J. Mentzer, P. Supasa, H. M. E. Duyvesteyn, R. Das, D. Skelly, T. G. Ritter, A. Amini, S. Bibi, S. Adele, S. A. Johnson, B. Constantinides, H. Webster, N. Temperton, P. Klenerman, E. Barnes, S. J. Dunachie, D. Crook, A. J. Pollard, T. Lambe, P. Goulder, N. G. Paterson, M. A. Williams, D. R. Hall, C. Conlon, A. Deeks, J. Frater, L. Frending, S. Gardiner, A. Jämsén, K. Jeffery, T. Malone, E. Phillips, L. Rothwell, L. Stafford, E. E. Fry, J. Huo, J. Mongkolsapaya, J. Ren, D. I. Stuart, G. R. Screaton, Antibody escape of SARS-CoV-2 Omicron BA.4 and BA.5 from vaccine and BA.1 serum. Cell 185, 2422-2433.e13 (2022).

4. R. Viana, S. Moyo, D. G. Amoako, H. Tegally, C. Scheepers, C. L. Althaus, U. J. Anyaneji, P. A. Bester, M. F. Boni, M. Chand, W. T. Choga, R. Colquhoun, M. Davids, K. Deforche, D. Doolabh, L. du Plessis, S. Engelbrecht, J. Everatt, J. Giandhari, M. Giovanetti, D. Hardie, V. Hill, N.-Y. Hsiao, A. Iranzadeh, A. Ismail, C. Joseph, R. Joseph, L. Koopile, S. L. Kosakovsky Pond, M. U. G. Kraemer, L. Kuate-Lere, O. Laguda-Akingba, O. Lesetedi-Mafoko, R. J. Lessells, S. Lockman, A. G. Lucaci, A. Maharaj, B. Mahlangu, T. Maponga, K. Mahlakwane, Z. Makatini, G. Marais, D. Maruapula, K. Masupu, M. Matshaba, S. Mayaphi, N. Mbhele, M. B. Mbulawa, A. Mendes, K. Mlisana, A. Mnguni, T. Mohale, M. Moir, K. Moruisi, M. Mosepele, G. Motsatsi, M. S. Motswaledi, T. Mphoyakgosi, N. Msomi, P. N. Mwangi, Y. Naidoo, N. Ntuli, M. Nyaga, L. Olubayo, S. Pillay, B. Radibe, Y. Ramphal, U. Ramphal, J. E. San, L. Scott, R. Shapiro, L. Singh, P. Smith-Lawrence, W. Stevens, A. Strydom, K. Subramoney, N. Tebeila, D. Tshiabuila, J. Tsui, S. van Wyk, S. Weaver, C. K. Wibmer, E. Wilkinson, N. Wolter, A. E. Zarebski, B. Zuze, D. Goedhals, W. Preiser, F. Treurnicht, M. Venter, C. Williamson, O. G. Pybus, J. Bhiman, A. Glass, D. P. Martin, A. Rambaut, S. Gaseitsiwe, A. von Gottberg, T. de Oliveira, Rapid epidemic expansion of the SARS-CoV-2 Omicron variant in southern Africa. Nature 603, 679–686 (2022).

5. Q. Wang, S. Iketani, Z. Li, L. Liu, Y. Guo, Y. Huang, A. D. Bowen, M. Liu, M. Wang, J. Yu, R. Valdez, A. S. Lauring, Z. Sheng, H. H. Wang, A. Gordon, L. Liu, D. D. Ho, Alarming antibody evasion properties of rising SARS-CoV-2 BQ and XBB subvariants. Cell 186, 279–286 (2022).

6. D. Mannar, J. W. Saville, X. Zhu, S. S. Srivastava, A. M. Berezuk, K. S. Tuttle, A. C. Marquez, I. Sekirov, S. Subramaniam, SARS-CoV-2 Omicron variant: Antibody evasion and cryo-EM structure of spike protein–ACE2 complex. Science 375, 760–764 (2022).

7. E. Callaway, Are COVID surges becoming more predictable? New Omicron variants offer a hint. Nature 605, 204–206 (2022).

8. E. Callaway, COVID ‘variant soup’ is making winter surges hard to predict. Nature 611, 213–214 (2022).

9. H. Marcotte, L. Hammarström, Q. Pan-Hammarström, Limited cross-variant neutralization after primary Omicron infection: consideration for a variant-containing booster. Signal Transduct. Target. Ther. 7, 1–3 (2022).

10. E. Callaway, What Omicron’s BA.4 and BA.5 variants mean for the pandemic. Nature 606, 848–849 (2022).

11. S. Iketani, L. Liu, Y. Guo, L. Liu, J. F.-W. Chan, Y. Huang, M. Wang, Y. Luo, J. Yu, H. Chu, K. K.-H. Chik, T. T.-T. Yuen, M. T. Yin, M. E. Sobieszczyk, Y. Huang, K.-Y. Yuen, H. H. Wang, Z. Sheng, D. D. Ho, Antibody evasion properties of SARS-CoV-2 Omicron sublineages. Nature 604, 553–556 (2022).

12. Y. Cao, F. Jian, J. Wang, Y. Yu, W. Song, A. Yisimayi, J. Wang, R. An, X. Chen, N. Zhang, Y. Wang, P. Wang, L. Zhao, H. Sun, L. Yu, S. Yang, X. Niu, T. Xiao, Q. Gu, F. Shao, X. Hao, Y. Xu, R. Jin, Z. Shen, Y. Wang, X. S. Xie, Imprinted SARS-CoV-2 humoral immunity induces convergent Omicron RBD evolution. Nature (2022), doi: 10.1038/s41586-022-05644-7.

13. C. Hirsch, Y. S. Park, V. Piechotta, K. L. Chai, L. J. Estcourt, I. Monsef, S. Salomon, E. M. Wood, C. So-Osman, Z. McQuilten, C. D. Spinner, J. J. Malin, M. Stegemann, N. Skoetz, N. Kreuzberger, SARS-CoV-2-neutralising monoclonal antibodies to prevent COVID-19. Cochrane Database Syst. Rev. 6, CD014945 (2022).

14. L. Hammarström, H. Abolhassani, F. Baldanti, H. Marcotte, Q. Pan-Hammarström, Development of passive immunity against SARS-CoV-2 for management of immunodeficient patients—a perspective. J. Allergy Clin. Immunol. 146, 58–60 (2020).

15. L. Hammarström, H. Marcotte, A. Piralla, F. Baldanti, Q. Pan-Hammarström, Antibody therapy for COVID-19. Curr. Opin. Allergy Clin. Immunol. 21, 553–558 (2021).

16. Y. Nguyen, A. Flahault, N. Chavarot, C. Melenotte, M. Cheminant, P. Deschamps, N. Carlier, E. Lafont, M. Thomas, E. Flamarion, D. Lebeaux, C. Charlier, A. Rachline, C. Guérin, R. Ratiney, J. Touchard, H. Péré, F. Rozenberg, F. Lanternier, J.-B. Arlet, J. Avouac, V. Boussaud, R. Guillemain, M. Vignon, E. Thervet, A. Scemla, L. Weiss, L. Mouthon, AP-HP-Centre Monoclonal Antibodies Working Group, Pre-exposure prophylaxis with tixagevimab and cilgavimab (Evusheld) for COVID-19 among 1112 severely immunocompromised patients. Clin. Microbiol. Infect. 28, 1654.e1-1654.e4 (2022).

17. R. Uraki, P. J. Halfmann, S. Iida, S. Yamayoshi, Y. Furusawa, M. Kiso, M. Ito, K. Iwatsuki-Horimoto, S. Mine, M. Kuroda, T. Maemura, Y. Sakai-Tagawa, H. Ueki, R. Li, Y. Liu, D. Larson, S. Fukushi, S. Watanabe, K. Maeda, A. Pekosz, A. Kandeil, R. J. Webby, Z. Wang, M. Imai, T. Suzuki, Y. Kawaoka, Characterization of SARS-CoV-2 Omicron BA.4 and BA.5 isolates in rodents. Nature 612, 1–6 (2022).

18. M. Kozlov, Omicron’s feeble attack on the lungs could make it less dangerous. Nature 601, 177–177 (2022).

19. M. W. Russell, Z. Moldoveanu, P. L. Ogra, J. Mestecky, Mucosal Immunity in COVID-19: A Neglected but Critical Aspect of SARS-CoV-2 Infection. Front. Immunol. 11, 611337 (2020).

20. E. Waltz, How nasal-spray vaccines could change the pandemic. Nature 609, 240–242 (2022).

21. S. Kumar Bharathkar, B. W. Parker, A. G. Malyutin, N. Haloi, K. E. Huey-Tubman, E. Tajkhorshid, B. M. Stadtmueller, S. H. Scheres, J. Kuriyan, S. H. Scheres, M. F. Flajnik, Eds. The structures of secretory and dimeric immunoglobulin A. eLife 9, e56098 (2020).

22. P. Brandtzaeg, H. Prydz, Direct evidence for an integrated function of J chain and secretory component in epithelial transport of immunoglobulins. Nature 311, 71–73 (1984).

23. J. M. Woof, M. W. Russell, Structure and function relationships in IgA. Mucosal Immunol. 4, 590–597 (2011).

24. K. Kett, P. Brandtzaeg, J. Radl, J. J. Haaijman, Different subclass distribution of IgA-producing cells in human lymphoid organs and various secretory tissues. J. Immunol. 136, 3631–3635 (1986).

25. M. Lin, L. Du, P. Brandtzaeg, Q. Pan-Hammarström, IgA subclass switch recombination in human mucosal and systemic immune compartments. Mucosal Immunol. 7, 511–520 (2014).

26. D. Sterlin, A. Mathian, M. Miyara, A. Mohr, F. Anna, L. Claër, P. Quentric, J. Fadlallah, H. Devilliers, P. Ghillani, C. Gunn, R. Hockett, S. Mudumba, A. Guihot, C.-E. Luyt, J. Mayaux, A. Beurton, S. Fourati, T. Bruel, O. Schwartz, J.-M. Lacorte, H. Yssel, C. Parizot, K. Dorgham, P. Charneau, Z. Amoura, G. Gorochov, IgA dominates the early neutralizing antibody response to SARS-CoV-2. Sci. Transl. Med. 13, eabd2223 (2021).

27. S. Havervall, U. Marking, J. Svensson, N. Greilert-Norin, P. Bacchus, P. Nilsson, S. Hober, M. Gordon, K. Blom, J. Klingström, M. Åberg, A. Smed-Sörensen, C. Thålin, Anti-Spike Mucosal IgA Protection against SARS-CoV-2 Omicron Infection. N. Engl. J. Med. 387, 1333–1336 (2022).

28. Z. Wang, J. C. C. Lorenzi, F. Muecksch, S. Finkin, C. Viant, C. Gaebler, M. Cipolla, H.-H. Hoffmann, T. Y. Oliveira, D. A. Oren, V. Ramos, L. Nogueira, E. Michailidis, D. F. Robbiani, A. Gazumyan, C. M. Rice, T. Hatziioannou, P. D. Bieniasz, M. Caskey, M. C. Nussenzweig, Enhanced SARS-CoV-2 neutralization by dimeric IgA. Sci. Transl. Med. 13, eabf1555 (2021).

29. B. Isho, K. T. Abe, M. Zuo, A. J. Jamal, B. Rathod, J. H. Wang, Z. Li, G. Chao, O. L. Rojas, Y. M. Bang, A. Pu, N. Christie-Holmes, C. Gervais, D. Ceccarelli, P. Samavarchi-Tehrani, F. Guvenc, P. Budylowski, A. Li, A. Paterson, F. Y. Yue, L. M. Marin, L. Caldwell, J. L. Wrana, K. Colwill, F. Sicheri, S. Mubareka, S. D. Gray-Owen, S. J. Drews, W. L. Siqueira, M. Barrios-Rodiles, M. Ostrowski, J. M. Rini, Y. Durocher, A. J. McGeer, J. L. Gommerman, A.-C. Gingras, Persistence of serum and saliva antibody responses to SARS-CoV-2 spike antigens in COVID-19 patients. Sci. Immunol. 5, eabe5511 (2020).

30. N. Huang, P. Pérez, T. Kato, Y. Mikami, K. Okuda, R. C. Gilmore, C. D. Conde, B. Gasmi, S. Stein, M. Beach, E. Pelayo, J. O. Maldonado, B. A. Lafont, S.-I. Jang, N. Nasir, R. J. Padilla, V. A. Murrah, R. Maile, W. Lovell, S. M. Wallet, N. M. Bowman, S. L. Meinig, M. C. Wolfgang, S. N. Choudhury, M. Novotny, B. D. Aevermann, R. H. Scheuermann, G. Cannon, C. W. Anderson, R. E. Lee, J. T. Marchesan, M. Bush, M. Freire, A. J. Kimple, D. L. Herr, J. Rabin, A. Grazioli, S. Das, B. N. French, T. Pranzatelli, J. A. Chiorini, D. E. Kleiner, S. Pittaluga, S. M. Hewitt, P. D. Burbelo, D. Chertow, K. Frank, J. Lee, R. C. Boucher, S. A. Teichmann, B. M. Warner, K. M. Byrd, SARS-CoV-2 infection of the oral cavity and saliva. Nat. Med. 27, 892–903 (2021).

31. Zuo, F., Marcotte, H., Hammarström, L., Pan-Hammarström, Q., Mucosal IgA against SARS-CoV-2 Omicron Infection. N. Engl. J. Med. 387, e55 (2022).

32. Q. Yan, P. He, X. Huang, K. Luo, Y. Zhang, H. Yi, Q. Wang, F. Li, R. Hou, X. Fan, P. Li, X. Liu, H. Liang, Y. Deng, Z. Chen, Y. Chen, X. Mo, L. Feng, X. Xiong, S. Li, J. Han, L. Qu, X. Niu, L. Chen, Germline IGHV3-53-encoded RBD-targeting neutralizing antibodies are commonly present in the antibody repertoires of COVID-19 patients. Emerg. Microbes Infect. 10, 1097–1111.

33. S. Du, Y. Cao, Q. Zhu, P. Yu, F. Qi, G. Wang, X. Du, L. Bao, W. Deng, H. Zhu, J. Liu, J. Nie, Y. Zheng, H. Liang, R. Liu, S. Gong, H. Xu, A. Yisimayi, Q. Lv, B. Wang, R. He, Y. Han, W. Zhao, Y. Bai, Y. Qu, X. Gao, C. Ji, Q. Wang, N. Gao, W. Huang, Y. Wang, X. S. Xie, X. Su, J. Xiao, C. Qin, Structurally Resolved SARS-CoV-2 Antibody Shows High Efficacy in Severely Infected Hamsters and Provides a Potent Cocktail Pairing Strategy. Cell 183, 1013–1023.e13 (2020).

34. Y. Zhou, Z. Liu, S. Li, W. Xu, Q. Zhang, I. T. Silva, C. Li, Y. Wu, Q. Jiang, Z. Liu, Q. Wang, Y. Guo, J. Wu, C. Gu, X. Cai, D. Qu, C. T. Mayer, X. Wang, S. Jiang, T. Ying, Z. Yuan, Y. Xie, Y. Wen, L. Lu, Q. Wang, Enhancement versus neutralization by SARS-CoV-2 antibodies from a convalescent donor associates with distinct epitopes on the RBD. Cell Rep. 34, 108699 (2021).

35. Z. Liu, W. Xu, Z. Chen, W. Fu, W. Zhan, Y. Gao, J. Zhou, Y. Zhou, J. Wu, Q. Wang, X. Zhang, A. Hao, W. Wu, Q. Zhang, Y. Li, K. Fan, R. Chen, Q. Jiang, C. T. Mayer, T. Schoofs, Y. Xie, S. Jiang, Y. Wen, Z. Yuan, K. Wang, L. Lu, L. Sun, Q. Wang, An ultrapotent pan-β-coronavirus lineage B (β-CoV-B) neutralizing antibody locks the receptor-binding domain in closed conformation by targeting its conserved epitope. Protein Cell 13, 655–675 (2021).

36. C. O. Barnes, C. A. Jette, M. E. Abernathy, K.-M. A. Dam, S. R. Esswein, H. B. Gristick, A. G. Malyutin, N. G. Sharaf, K. E. Huey-Tubman, Y. E. Lee, D. F. Robbiani, M. C. Nussenzweig, A. P. West, P. J. Bjorkman, SARS-CoV-2 neutralizing antibody structures inform therapeutic strategies. Nature 588, 682–687 (2020).

37. L. Liu, S. Iketani, Y. Guo, J. F.-W. Chan, M. Wang, L. Liu, Y. Luo, H. Chu, Y. Huang, M. S. Nair, J. Yu, K. K.-H. Chik, T. T.-T. Yuen, C. Yoon, K. K.-W. To, H. Chen, M. T. Yin, M. E. Sobieszczyk, Y. Huang, H. H. Wang, Z. Sheng, K.-Y. Yuen, D. D. Ho, Striking antibody evasion manifested by the Omicron variant of SARS-CoV-2. Nature 602, 676–681 (2022).

38. M. F. Boni, P. Lemey, X. Jiang, T. T.-Y. Lam, B. W. Perry, T. A. Castoe, A. Rambaut, D. L. Robertson, Evolutionary origins of the SARS-CoV-2 sarbecovirus lineage responsible for the COVID-19 pandemic. Nat. Microbiol. 5, 1408–1417 (2020).

39. T. N. Starr, A. J. Greaney, S. K. Hilton, D. Ellis, K. H. D. Crawford, A. S. Dingens, M. J. Navarro, J. E. Bowen, M. A. Tortorici, A. C. Walls, N. P. King, D. Veesler, J. D. Bloom, Deep Mutational Scanning of SARS-CoV-2 Receptor Binding Domain Reveals Constraints on Folding and ACE2 Binding. Cell 182, 1295–1310.e20 (2020).

40. Z. Ku, X. Xie, P. R. Hinton, X. Liu, X. Ye, A. E. Muruato, D. C. Ng, S. Biswas, J. Zou, Y. Liu, D. Pandya, V. D. Menachery, S. Rahman, Y.-A. Cao, H. Deng, W. Xiong, K. B. Carlin, J. Liu, H. Su, E. J. Haanes, B. A. Keyt, N. Zhang, S. F. Carroll, P.-Y. Shi, Z. An, Nasal delivery of an IgM offers broad protection from SARS-CoV-2 variants. Nature 595, 718–723 (2021).

41. R. De Gasparo, M. Pedotti, L. Simonelli, P. Nickl, F. Muecksch, I. Cassaniti, E. Percivalle, J. C. C. Lorenzi, F. Mazzola, D. Magrì, T. Michalcikova, J. Haviernik, V. Honig, B. Mrazkova, N. Polakova, A. Fortova, J. Tureckova, V. Iatsiuk, S. Di Girolamo, M. Palus, D. Zudova, P. Bednar, I. Bukova, F. Bianchini, D. Mehn, R. Nencka, P. Strakova, O. Pavlis, J. Rozman, S. Gioria, J. C. Sammartino, F. Giardina, S. Gaiarsa, Q. Pan-Hammarström, C. O. Barnes, P. J. Bjorkman, L. Calzolai, A. Piralla, F. Baldanti, M. C. Nussenzweig, P. D. Bieniasz, T. Hatziioannou, J. Prochazka, R. Sedlacek, D. F. Robbiani, D. Ruzek, L. Varani, Bispecific IgG neutralizes SARS-CoV-2 variants and prevents escape in mice. Nature 593, 424–428 (2021).

42. Z. Ke, J. Oton, K. Qu, M. Cortese, V. Zila, L. McKeane, T. Nakane, J. Zivanov, C. J. Neufeldt, B. Cerikan, J. M. Lu, J. Peukes, X. Xiong, H.-G. Kräusslich, S. H. W. Scheres, R. Bartenschlager, J. A. G. Briggs, Structures and distributions of SARS-CoV-2 spike proteins on intact virions. Nature 588, 498–502 (2020).

43. M. F. Bachmann, M. O. Mohsen, L. Zha, M. Vogel, D. E. Speiser, SARS-CoV-2 structural features may explain limited neutralizing-antibody responses. Npj Vaccines 6, 1–5 (2021).

44. P. Qu, J. P. Evans, Y.-M. Zheng, C. Carlin, L. J. Saif, E. M. Oltz, K. Xu, R. J. Gumina, S.-L. Liu, Evasion of neutralizing antibody responses by the SARS-CoV-2 BA.2.75 variant. Cell Host Microbe 30, 1518-1526.e4 (2022).

45. Q. Wang, Y. Guo, S. Iketani, M. S. Nair, Z. Li, H. Mohri, M. Wang, J. Yu, A. D. Bowen, J. Y. Chang, J. G. Shah, N. Nguyen, Z. Chen, K. Meyers, M. T. Yin, M. E. Sobieszczyk, Z. Sheng, Y. Huang, L. Liu, D. D. Ho, Antibody evasion by SARS-CoV-2 Omicron subvariants BA.2.12.1, BA.4 and BA.5. Nature 608, 603–608 (2022).

46. Y. Cao, F. Jian, Z. Zhang, A. Yisimayi, X. Hao, L. Bao, F. Yuan, Y. Yu, S. Du, J. Wang, T. Xiao, W. Song, Y. Zhang, P. Liu, R. An, P. Wang, Y. Wang, S. Yang, X. Niu, Y. Zhang, Q. Gu, F. Shao, Y. Hu, W. Yin, A. Zheng, Y. Wang, C. Qin, R. Jin, J. Xiao, X. S. Xie, Rational identification of potent and broad sarbecovirus-neutralizing antibody cocktails from SARS convalescents. Cell Rep. 41, 11185 (2022).

47. F. Bianchini, V. Crivelli, M. E. Abernathy, C. Guerra, M. Palus, J. Muri, H. Marcotte, A. Piralla, M. Pedotti, R. De Gasparo, L. Simonelli, M. Matkovic, C. Toscano, M. Biggiogero, V. Calvaruso, P. Svoboda, T. Cervantes Rincón, T. Fava, L. Podešvová, A. A. Shanbhag, A. Celoria, J. Sgrignani, M. Stefanik, V. Hönig, V. Pranclova, T. Michalcikova, J. Prochazka, G. Guerrini, D. Mehn, A. Ciabattini, H. Abolhassani, D. Jarrossay, M. Uguccioni, D. Medaglini, Q. Pan-Hammarström, L. Calzolai, D. Fernandez, F. Baldanti, A. Franzetti-Pellanda, C. Garzoni, R. Sedlacek, D. Ruzek, L. Varani, A. Cavalli, C. O. Barnes, D. F. Robbiani, Human neutralizing antibodies to cold linear epitopes and subdomain 1 of the SARS-CoV-2 spike glycoprotein. Sci. Immunol., eade0958 (2023).

48. F. Su, G. B. Patel, S. Hu, W. Chen, Induction of mucosal immunity through systemic immunization: Phantom or reality? Hum. Vaccines Immunother. 12, 1070–1079 (2016).

49. K. Sano, D. Bhavsar, G. Singh, D. Floda, K. Srivastava, C. Gleason, J. M. Carreño, V. Simon, F. Krammer, SARS-CoV-2 vaccination induces mucosal antibody responses in previously infected individuals. Nat. Commun. 13, 5135 (2022).

50. H. M. Joo, Y. He, A. Sundararajan, L. Huan, M. Y. Sangster, Quantitative analysis of influenza virus-specific B cell memory generated by different routes of inactivated virus vaccination. Vaccine 28, 2186–2194 (2010).

51. T. Mao, B. Israelow, M. A. Peña-Hernández, A. Suberi, L. Zhou, S. Luyten, M. Reschke, H. Dong, R. J. Homer, W. M. Saltzman, A. Iwasaki, Unadjuvanted intranasal spike vaccine elicits protective mucosal immunity against sarbecoviruses. Science 378, eabo2523 (2022).

52. Y. Cao, A. Yisimayi, F. Jian, W. Song, T. Xiao, L. Wang, S. Du, J. Wang, Q. Li, X. Chen, Y. Yu, P. Wang, Z. Zhang, P. Liu, R. An, X. Hao, Y. Wang, J. Wang, R. Feng, H. Sun, L. Zhao, W. Zhang, D. Zhao, J. Zheng, L. Yu, C. Li, N. Zhang, R. Wang, X. Niu, S. Yang, X. Song, Y. Chai, Y. Hu, Y. Shi, L. Zheng, Z. Li, Q. Gu, F. Shao, W. Huang, R. Jin, Z. Shen, Y. Wang, X. Wang, J. Xiao, X. S. Xie, BA.2.12.1, BA.4 and BA.5 escape antibodies elicited by Omicron infection. Nature 608, 593–602 (2022).

53. K. Westendorf, S. Žentelis, L. Wang, D. Foster, P. Vaillancourt, M. Wiggin, E. Lovett, R. van der Lee, J. Hendle, A. Pustilnik, J. M. Sauder, L. Kraft, Y. Hwang, R. W. Siegel, J. Chen, B. A. Heinz, R. E. Higgs, N. L. Kallewaard, K. Jepson, R. Goya, M. A. Smith, D. W. Collins, D. Pellacani, P. Xiang, V. de Puyraimond, M. Ricicova, L. Devorkin, C. Pritchard, A. O’Neill, K. Dalal, P. Panwar, H. Dhupar, F. A. Garces, C. A. Cohen, J. M. Dye, K. E. Huie, C. V. Badger, D. Kobasa, J. Audet, J. J. Freitas, S. Hassanali, I. Hughes, L. Munoz, H. C. Palma, B. Ramamurthy, R. W. Cross, T. W. Geisbert, V. Menachery, K. Lokugamage, V. Borisevich, I. Lanz, L. Anderson, P. Sipahimalani, K. S. Corbett, E. S. Yang, Y. Zhang, W. Shi, T. Zhou, M. Choe, J. Misasi, P. D. Kwong, N. J. Sullivan, B. S. Graham, T. L. Fernandez, C. L. Hansen, E. Falconer, J. R. Mascola, B. E. Jones, B. C. Barnhart, LY-CoV1404 (bebtelovimab) potently neutralizes SARS-CoV-2 variants. Cell Rep. 39, 110812 (2022).

54. P. Turelli, C. Fenwick, C. Raclot, V. Genet, G. Pantaleo, D. Trono, P2G3 human monoclonal antibody neutralizes SARS-CoV-2 Omicron subvariants including BA.4 and BA.5 and Bebtelovimab escape mutants. bioRxiv (2022), https://doi.org/10.1101/2022.07.28.501852.

55. J. B. Case, S. Mackin, J. M. Errico, Z. Chong, E. A. Madden, B. Whitener, B. Guarino, M. A. Schmid, K. Rosenthal, K. Ren, H. V. Dang, G. Snell, A. Jung, L. Droit, S. A. Handley, P. J. Halfmann, Y. Kawaoka, J. E. Crowe, D. H. Fremont, H. W. Virgin, Y.-M. Loo, M. T. Esser, L. A. Purcell, D. Corti, M. S. Diamond, Resilience of S309 and AZD7442 monoclonal antibody treatments against infection by SARS-CoV-2 Omicron lineage strains. Nat. Commun. 13, 3824 (2022).

56. X. Li, Y. Pan, Q. Yin, Z. Wang, S. Shan, L. Zhang, J. Yu, Y. Qu, L. Sun, F. Gui, J. Lu, Z. Jing, W. Wu, T. Huang, X. Shi, J. Li, X. Li, D. Li, S. Wang, M. Yang, L. Zhang, K. Duan, M. Liang, X. Yang, X. Wang, Structural basis of a two-antibody cocktail exhibiting highly potent and broadly neutralizing activities against SARS-CoV-2 variants including diverse Omicron sublineages. Cell Discov. 8, 87 (2022).

57. H. Gruell, K. Vanshylla, M. Korenkov, P. Tober-Lau, M. Zehner, F. Münn, H. Janicki, M. Augustin, P. Schommers, L. E. Sander, F. Kurth, C. Kreer, F. Klein, SARS-CoV-2 Omicron sublineages exhibit distinct antibody escape patterns. Cell Host Microbe 30, 1–11 (2022).

58. S. Luo, J. Zhang, A. J. B. Kreutzberger, A. Eaton, R. J. Edwards, C. Jing, H.-Q. Dai, G. D. Sempowski, K. Cronin, R. Parks, A. Y. Ye, K. Mansouri, M. Barr, N. Pishesha, A. C. Williams, L. Vieira Francisco, A. Saminathan, H. Peng, H. Batra, L. Bellusci, S. Khurana, S. M. Alam, D. C. Montefiori, K. O. Saunders, M. Tian, H. Ploegh, T. Kirchhausen, B. Chen, B. F. Haynes, F. W. Alt, An antibody from single human VH-rearranging mouse neutralizes all SARS-CoV-2 variants through BA.5 by inhibiting membrane fusion. Sci. Immunol. 7, eadd5446 (2022).

59. A. Bonner, P. B. Furtado, A. Almogren, M. A. Kerr, S. J. Perkins, Implications of the Near-Planar Solution Structure of Human Myeloma Dimeric IgA1 for Mucosal Immunity and IgA Nephropathy. J. Immunol. 180, 1008–1018 (2008).

60. A. Bonner, A. Almogren, P. B. Furtado, M. A. Kerr, S. J. Perkins, The Nonplanar Secretory IgA2 and Near Planar Secretory IgA1 Solution Structures Rationalize Their Different Mucosal Immune Responses. J. Biol. Chem. 284, 5077–5087 (2009).

61. M. Ejemel, Q. Li, S. Hou, Z. A. Schiller, J. A. Tree, A. Wallace, A. Amcheslavsky, N. Kurt Yilmaz, K. R. Buttigieg, M. J. Elmore, K. Godwin, N. Coombes, J. R. Toomey, R. Schneider, A. S. Ramchetty, B. J. Close, D.-Y. Chen, H. L. Conway, M. Saeed, C. Ganesa, M. W. Carroll, L. A. Cavacini, M. S. Klempner, C. A. Schiffer, Y. Wang, A cross-reactive human IgA monoclonal antibody blocks SARS-CoV-2 spike-ACE2 interaction. Nat. Commun. 11, 4198 (2020).

62. A. Natarajan, S. Zlitni, E. F. Brooks, S. E. Vance, A. Dahlen, H. Hedlin, R. M. Park, A. Han, D. T. Schmidtke, R. Verma, K. B. Jacobson, J. Parsonnet, H. F. Bonilla, U. Singh, B. A. Pinsky, J. R. Andrews, P. Jagannathan, A. S. Bhatt, Gastrointestinal symptoms and fecal shedding of SARS-CoV-2 RNA suggest prolonged gastrointestinal infection. Med 3, 371–387.e9 (2022).

63. Y. Wang, A. Yan, D. Song, C. Dong, M. Rao, Y. Gao, R. Qi, X. Ma, Q. Wang, H. Xu, H. Liu, J. Han, M. Duan, S. Liu, X. Yu, M. Zong, J. Feng, J. Jiao, H. Zhang, M. Li, B. Yu, Y. Wang, F. Meng, X. Ni, Y. Li, Z. Shen, B. Sun, X. Shao, H. Zhao, Y. Zhao, R. Li, Y. Zhang, G. Du, J. Lu, C. You, H. Jiang, L. Zhang, L. Wang, C. Dou, Z. Liu, J. Zhao, Biparatopic antibody BA7208/7125 effectively neutralizes SARS-CoV-2 variants including Omicron BA.1-BA.5. Cell Discov. 9, 1–16 (2023).

64. A. Y.-H. Teh, L. Cavacini, Y. Hu, O. S. Kumru, J. Xiong, D. T. Bolick, S. B. Joshi, C. Grünwald-Gruber, F. Altmann, M. Klempner, R. L. Guerrant, D. B. Volkin, Y. Wang, J. K.-C. Ma, Investigation of a monoclonal antibody against enterotoxigenic Escherichia coli, expressed as secretory IgA1 and IgA2 in plants. Gut Microbes 13, 1859813 (2021).

65. H. Marcotte, A. Piralla, F. Zuo, L. Du, I. Cassaniti, H. Wan, M. Kumagai-Braesh, J. Andréll, E. Percivalle, J. C. Sammartino, Y. Wang, S. Vlachiotis, J. Attevall, F. Bergami, A. Ferrari, M. Colaneri, M. Vecchia, M. Sambo, V. Zuccaro, E. Asperges, R. Bruno, T. Oggionni, F. Meloni, H. Abolhassani, F. Bertoglio, M. Schubert, L. Calzolai, L. Varani, M. Hust, Y. Xue, L. Hammarström, F. Baldanti, Q. Pan-Hammarström, Immunity to SARS-CoV-2 up to 15 months after infection. iScience 25, 103743 (2022).

66. N. Sherina, A. Piralla, L. Du, H. Wan, M. Kumagai-Braesch, J. Andréll, S. Braesch-Andersen, I. Cassaniti, E. Percivalle, A. Sarasini, F. Bergami, R. Di Martino, M. Colaneri, M. Vecchia, M. Sambo, V. Zuccaro, R. Bruno, M. Sachs, T. Oggionni, F. Meloni, H. Abolhassani, F. Bertoglio, M. Schubert, M. Byrne-Steele, J. Han, M. Hust, Y. Xue, L. Hammarström, F. Baldanti, H. Marcotte, Q. Pan-Hammarström, Persistence of SARS-CoV-2-specific B and T cell responses in convalescent COVID-19 patients 6–8 months after the infection. Med 2, 281–295.e4 (2021).

67. F. Bertoglio, V. Fühner, M. Ruschig, P. A. Heine, L. Abassi, T. Klünemann, U. Rand, D. Meier, N. Langreder, S. Steinke, R. Ballmann, K.-T. Schneider, K. D. R. Roth, P. Kuhn, P. Riese, D. Schäckermann, J. Korn, A. Koch, M. Z. Chaudhry, K. Eschke, Y. Kim, S. Zock-Emmenthal, M. Becker, M. Scholz, G. M. S. G. Moreira, E. V. Wenzel, G. Russo, H. S. P. Garritsen, S. Casu, A. Gerstner, G. Roth, J. Adler, J. Trimpert, A. Hermann, T. Schirrmann, S. Dübel, A. Frenzel, J. Van den Heuvel, L. Čičin-Šain, M. Schubert, M. Hust, A SARS-CoV-2 neutralizing antibody selected from COVID-19 patients binds to the ACE2-RBD interface and is tolerant to most known RBD mutations. Cell Rep. 36, 109433 (2021).

68. J. Korn, D. Schäckermann, T. Kirmann, F. Bertoglio, S. Steinke, J. Heisig, M. Ruschig, G. Rojas, N. Langreder, E. V. Wenzel, K. D. R. Roth, M. Becker, D. Meier, J. van den Heuvel, M. Hust, S. Dübel, M. Schubert, Baculovirus-free insect cell expression system for high yield antibody and antigen production. Sci. Rep. 10, 21393 (2020).

69. F. Zuo, H. Abolhassani, L. Du, A. Piralla, F. Bertoglio, L. de Campos-Mata, H. Wan, M. Schubert, I. Cassaniti, Y. Wang, J. C. Sammartino, R. Sun, S. Vlachiotis, F. Bergami, M. Kumagai-Braesch, J. Andréll, Z. Zhang, Y. Xue, E. V. Wenzel, L. Calzolai, L. Varani, N. Rezaei, Z. Chavoshzadeh, F. Baldanti, M. Hust, L. Hammarström, H. Marcotte, Q. Pan-Hammarström, Heterologous immunization with inactivated vaccine followed by mRNA-booster elicits strong immunity against SARS-CoV-2 Omicron variant. Nat. Commun. 13, 2670 (2022).

70. F. Schmidt, Y. Weisblum, F. Muecksch, H.-H. Hoffmann, E. Michailidis, J. C. C. Lorenzi, P. Mendoza, M. Rutkowska, E. Bednarski, C. Gaebler, M. Agudelo, A. Cho, Z. Wang, A. Gazumyan, M. Cipolla, M. Caskey, D. F. Robbiani, M. C. Nussenzweig, C. M. Rice, T. Hatziioannou, P. D. Bieniasz, Measuring SARS-CoV-2 neutralizing antibody activity using pseudotyped and chimeric virusesSARS-CoV-2 neutralizing antibody activity. J. Exp. Med. 217, e20201181 (2020).

71. E. Percivalle, G. Cambiè, I. Cassaniti, E. V. Nepita, R. Maserati, A. Ferrari, R. D. Martino, P. Isernia, F. Mojoli, R. Bruno, M. Tirani, D. Cereda, C. Nicora, M. Lombardo, F. Baldanti, Prevalence of SARS-CoV-2 specific neutralising antibodies in blood donors from the Lodi Red Zone in Lombardy, Italy, as at 06 April 2020. Eurosurveillance 25, 2001031 (2020).

72. A. Sircar, E. T. Kim, J. J. Gray, RosettaAntibody: antibody variable region homology modeling server. Nucleic Acids Res. 37, W474–W479 (2009).

73. M. Pedotti, L. Simonelli, E. Livoti, L. Varani, Computational Docking of Antibody-Antigen Complexes, Opportunities and Pitfalls Illustrated by Influenza Hemagglutinin. Int. J. Mol. Sci. 12, 226–251 (2011).

74. L. Simonelli, M. Beltramello, Z. Yudina, A. Macagno, L. Calzolai, L. Varani, Rapid Structural Characterization of Human Antibody–Antigen Complexes through Experimentally Validated Computational Docking. J. Mol. Biol. 396, 1491–1507 (2010).

75. D. Van Der Spoel, E. Lindahl, B. Hess, G. Groenhof, A. E. Mark, H. J. C. Berendsen, GROMACS: Fast, flexible, and free. J. Comput. Chem. 26, 1701–1718 (2005).

