## Supplementary material for "Conversion of monoclonal IgG to dimeric and secretory IgA restores neutralizing ability and prevents infection of Omicron lineages"

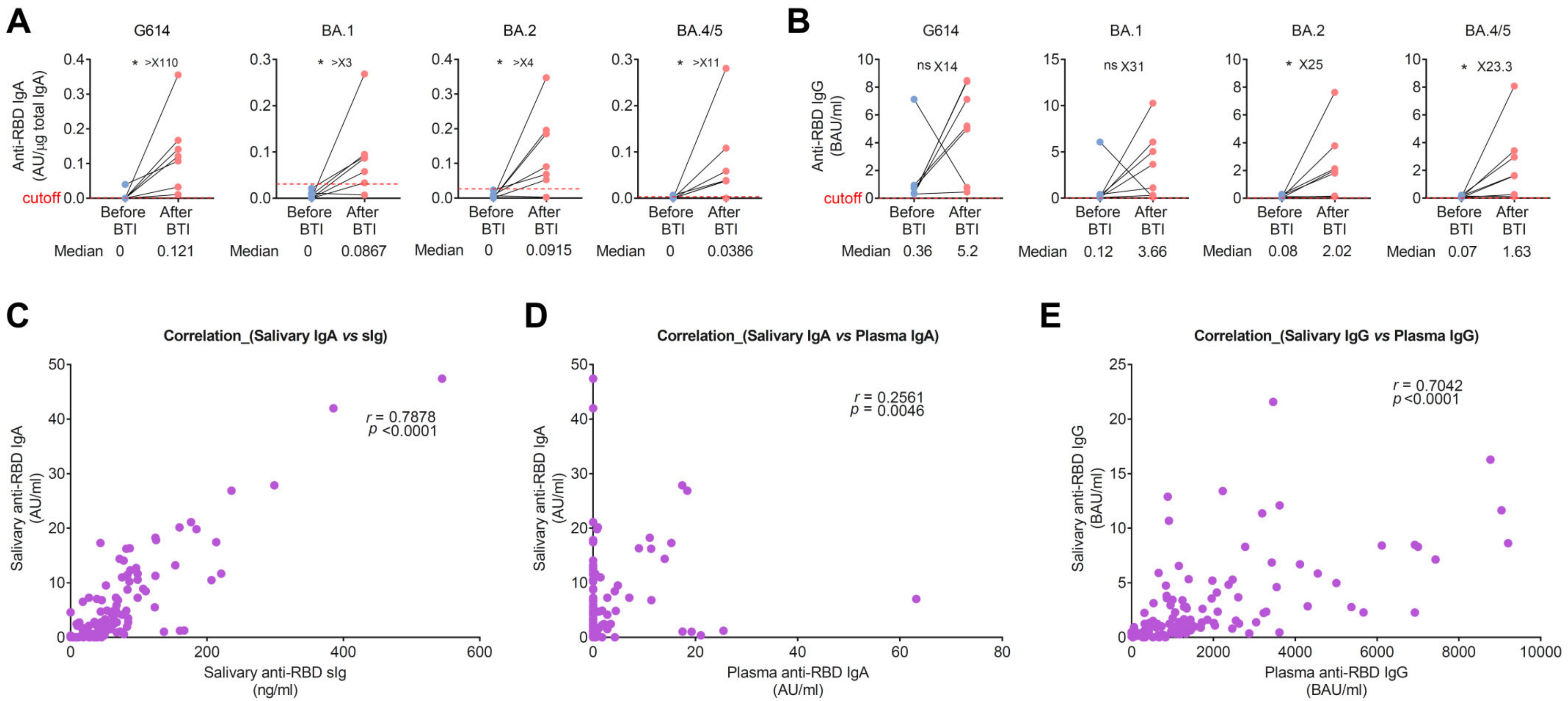


**Fig. S1. Salivary anti-RBD IgA antibodies correlate with salivary anti-RBD secretory immunoglobulins (sIg) and increase after breakthrough infection.** (**A** and **C**) Salivary anti-RBD IgA (A) and IgG (B) antibodies against G614 and Omicron variants BA.1, BA.2 and BA.4/5 in paired samples before and after breakthrough infection (BTI) in mRNA-vaccinated individuals. The number of fold differences in anti-RBD antibody titers are indicated. A Wilcoxon paired-sample signed-rank test was used. **P* < 0.05, and ***P* < 0.01. (**C** to **E**) Correlation between salivary anti-RBD IgA and salivary anti-RBD secretory immunoglobulin (sIg) (C), salivary anti-RBD IgA and plasma anti-RBD IgA (D), and salivary anti-RBD IgG and plasma anti-RBD IgG antibodies (E). Correlation analysis was performed using Spearman's rank correlation. Statistically significant if p < 0.05.

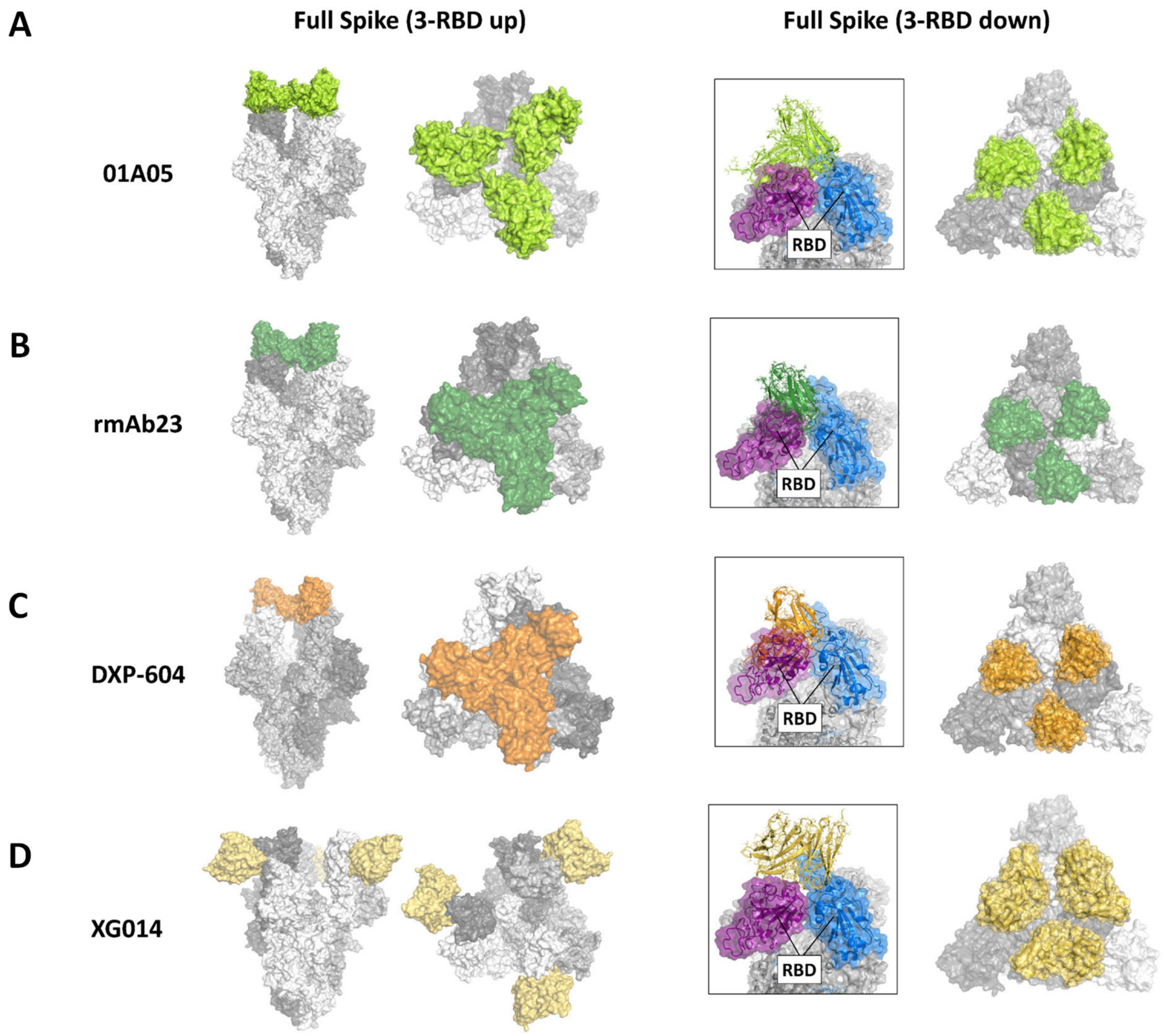


**Fig. S2. Computational simulations predicted the binding of neutralizing antibodies to RBDs in the S-trimer.** (**A** to **D**) Probability of 01A05 (A), rmAb23 (B), DXP-604 (C) and XG014 (D) binding to all three receptor-binding domains (RBDs) in the S-trimer (full S protein) in the up (3-RBD up) or down (3-RBD down) conformation. Three 01A05 can simultaneously bind the RBD on the S-trimer in the up conformation (3-RBD up) (A). One single rmAb23 Fab bound to the S-trimer with the 3-RBD in the up position (B). A single DXP-604 Fab bound to the S-trimer with the 3-RBD in the up position can prevent binding of ACE2 to all three S monomers and prevent the binding of other Fabs to S monomers (C). The epitopes of 01A05, mAb23 and DXP-604 are inaccessible on trimeric S 3-RBD down (A to C) because the antibodies interfere with the RBDs of the adjacent S monomer (in purple). Three XG014 Fabs can bind all three RBDs in the "down" conformation (3-RBD down) and should be able to bind the RBD in the up position (D).

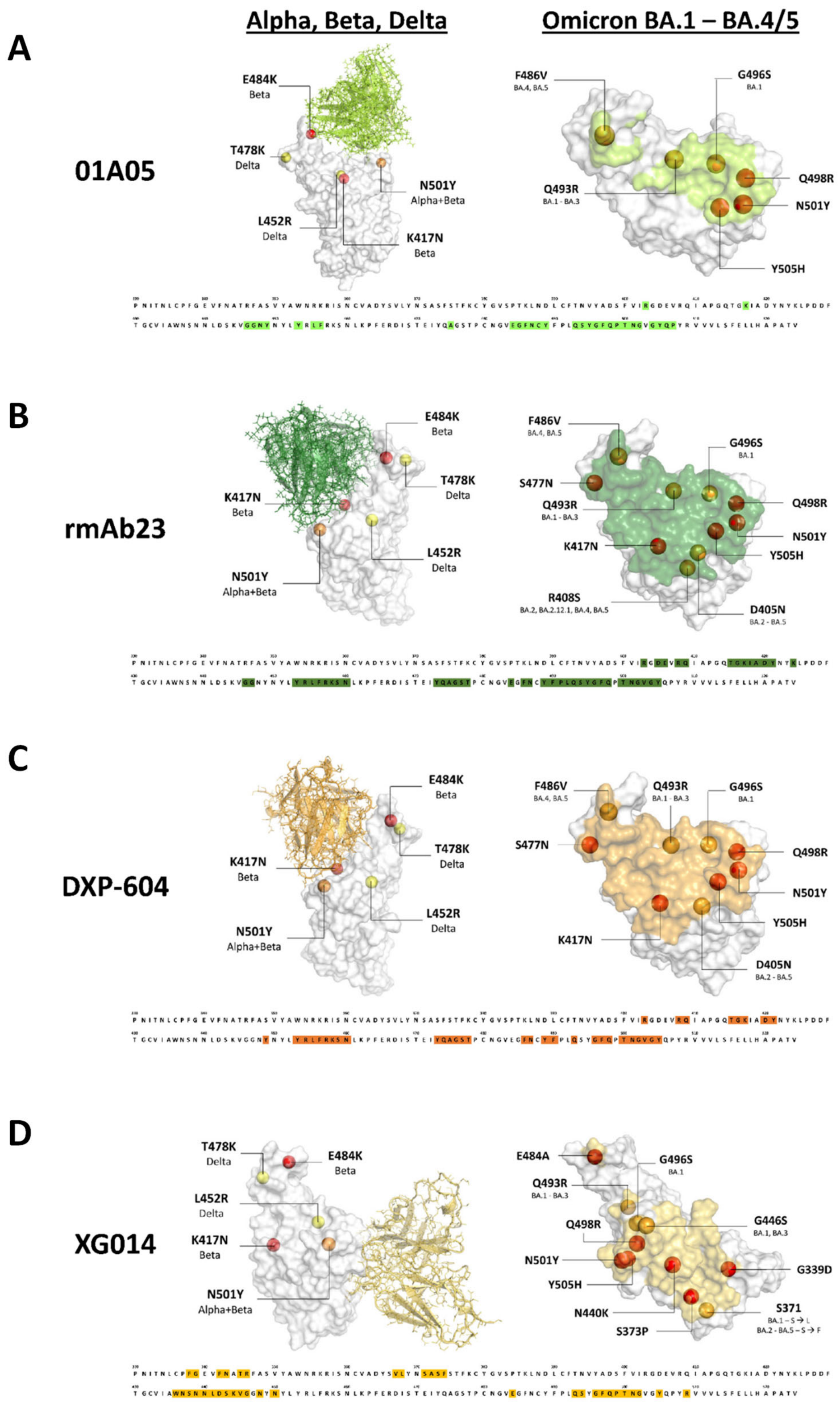


**Fig. S3. Computational simulations predicted antibody binding to the RBDs of variants of concern (VOCs).** (**A** to **D**) Docking model of 01A05 (A), rmAb23 (B), DXP-604 (C) and XG014 (D) to RBD of Alpha, Beta and Delta (left panel) and Omicron BA.1, BA.2 and BA.4/5 (right panel), including positions of mutations (red spheres). The sequence below each model shows the RBD epitope residues (highlighted) that make contact with an antibody.

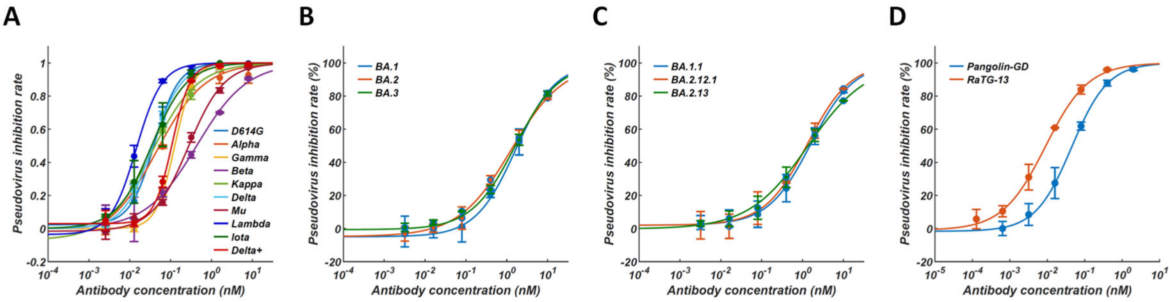


Pseudovirus neutralization IC50 (nM)

| Variants | Alpha | Beta | Gamma | Delta | Kappa | Delta plus | Mu | Lambda | Lota |
|---|---|---|---|---|---|---|---|---|---|
| | B.1.1.7 | B.1.351 | P.1 | B.1.617.2 | B.1.617.1 | AY.1 | B.1.621 | C.37 | B.1.621 |
| IC50 (nM) | 0.05 | 0.43 | 0.13 | 0.03 | 0.03 | 0.11 | 0.27 | 0.01 | 0.03 |

| Variants | Omicron | Omicron | Omicron | Omicron | Omicron | Omicron | RaTG13 | Pangolin |
|---|---|---|---|---|---|---|---|---|
| | BA.1 | BA.2 | BA.3 | BA.1.1 | BA.2.12.1 | BA.2.13 | | |
| IC50 (nM) | 1.53 | 1.2 | 1.6 | 1.6 | 1.33 | 1.33 | 0.01 | 0.05 |

IC50 (nM)

0-0.01

0.01-0.1

0.1-1

>1

**Fig. S4. DXP-604 broadly and potently neutralized SARS-CoV-2 variants. (A** to **D)** DXP-604 neutralization of SARS-CoV-2 variant S-pseudotyped VSV, including D614G, Alpha (B.1.1.7), Beta 453 (B.1.351), Gamma (P.1), Delta (B.1.617.2), 454 Kappa (B.1.617.1), Delta plus (AY.1), Mu (B.1.621), Lambda (C.37) and Iota (B.1.526) (A); BA.1, BA.2 and BA.3 (B); BA.1/BA.2 subvariants BA.1.1 (BA.1+R346K), BA.2.12.1 (BA.1+L452Q+S704L) and BA.2.13 (BA.1+L452M) (C); and clade 1b SARS-CoV-2 related sarbecoviruses (RaTG13 and Pangolin-GD) (D). The $IC_{50}$ values are indicated in the table.

## A 01A05

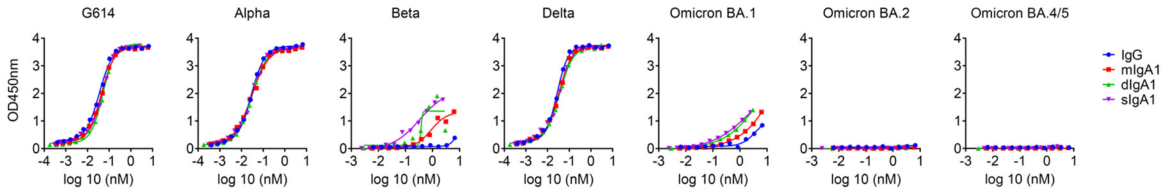


Binding EC50 (nM)

| | G614 | Alpha | Beta | Delta | Omicron BA.1 | Omicron BA.2 | Omicron BA.4/5 |
|---|---|---|---|---|---|---|---|
| IgG | 0.04 | 0.03 | >6.67 | 0.03 | >6.67 | >6.67 | >6.67 |
| mIgA1 | 0.04 | 0.03 | >6.25 | 0.03 | >6.25 | >6.25 | >6.25 |
| dIgA1 | 0.05 | 0.03 | >2.99 | 0.03 | >2.99 | >2.99 | >2.99 |
| sIgA1 | 0.05 | 0.03 | >2.44 | 0.03 | >2.44 | >2.44 | >2.44 |

EC50 (nM): 0-0.5, 0.5-2, 2-5, >5

Fold change to IgG

| | G614 | Alpha | Beta | Delta | Omicron BA.1 | Omicron BA.2 | Omicron BA.4/5 |
|---|---|---|---|---|---|---|---|
| mIgA1 | 0.82 | 0.96 | – | 0.25 | – | – | – |
| dIgA1 | 0.68 | 0.94 | – | 0.30 | – | – | – |
| sIgA1 | 0.73 | 0.95 | – | 0.26 | – | – | – |

Fold change: <1, 1-5, 5-10, 10-50, >50

### B rmAb23

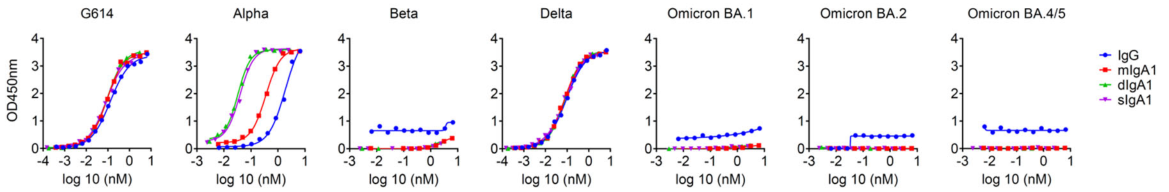


Binding EC50 (nM)

| | G614 | Alpha | Beta | Delta | Omicron BA.1 | Omicron BA.2 | Omicron BA.4/5 |
|---|---|---|---|---|---|---|---|
| IgG | 0.13 | 1.89 | >6.67 | 0.09 | >6.67 | >6.67 | >6.67 |
| mIgA1 | 0.09 | 0.35 | >6.25 | 0.07 | >6.25 | >6.25 | >6.25 |
| dIgA1 | 0.09 | 0.03 | >2.99 | 0.08 | >2.99 | >2.99 | >2.99 |
| sIgA1 | 0.08 | 0.04 | >2.44 | 0.08 | >2.44 | >2.44 | >2.44 |

EC50 (nM): 0-0.5, 0.5-2, 2-5, >5

Fold change to IgG

| | G614 | Alpha | Beta | Delta | Omicron BA.1 | Omicron BA.2 | Omicron BA.4/5 |
|---|---|---|---|---|---|---|---|
| mIgA1 | 1.42 | 5.38 | – | 1.25 | – | – | – |
| dIgA1 | 1.41 | 57.89 | – | 1.17 | – | – | – |
| sIgA1 | 1.53 | 48.58 | – | 1.13 | – | – | – |

Fold change: <1, 1-5, 5-10, 10-50, >50

## C XG014

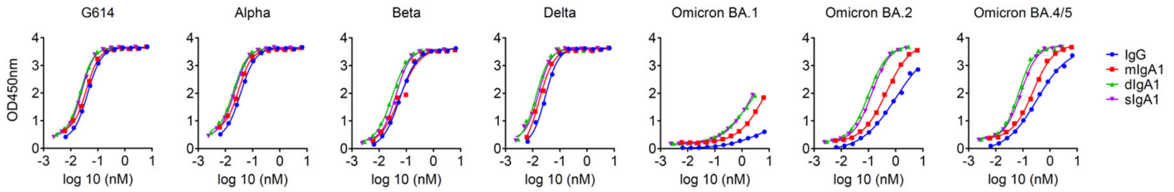


Binding EC50 (nM)

| | G614 | Alpha | Beta | Delta | Omicron BA.1 | Omicron BA.2 | Omicron BA.4/5 |
|---|---|---|---|---|---|---|---|
| IgG | 0.04 | 0.04 | 0.06 | 0.03 | 3.98 | 1.02 | 0.33 |
| mIgA1 | 0.04 | 0.03 | 0.06 | 0.02 | >6.25 | 0.45 | 0.24 |
| dIgA1 | 0.02 | 0.02 | 0.03 | 0.01 | 1.77 | 0.10 | 0.07 |
| sIgA1 | 0.03 | 0.02 | 0.04 | 0.02 | >2.44 | 0.13 | 0.09 |

EC50 (nM): 0-0.5, 0.5-2, 2-5, >5

Fold change to IgG

| | G614 | Alpha | Beta | Delta | Omicron BA.1 | Omicron BA.2 | Omicron BA.4/5 |
|---|---|---|---|---|---|---|---|
| mIgA1 | 1.12 | 1.17 | 0.99 | 1.65 | <0.63 | 2.25 | 1.41 |
| dIgA1 | 1.81 | 1.83 | 1.89 | 2.16 | 2.25 | 9.89 | 4.71 |
| sIgA1 | 1.66 | 1.67 | 1.39 | 1.75 | <1.63 | 7.71 | 3.88 |

Fold change: <1, 1-5, 5-10, 10-50, >50

**Fig. S5. Dimeric and secretory IgA1 enhanced binding activity against variants of concern (VOCs).** (**A** to **C**) Binding of IgG and IgA1 antibodies 01A05 (A), rmAb23 (B), and XG014 (C) to the RBD of G614 and VOCs (Alpha, Beta, Delta and Omicron), as determined by ELISAs. The $EC_{50}$ and fold-change differences between the IgG and IgA antibody forms are indicated.

## A 01A05

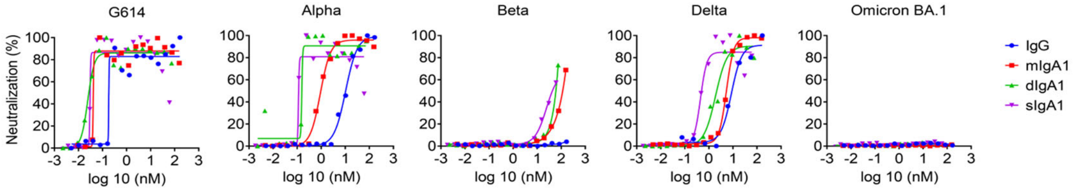


| Neutralization IC50 (nM) | G614 | Alpha | Beta | Delta | Omicron BA.1 |
|---|---|---|---|---|---|
| IgG | 0.18 | 10.41 | >167 | 8.62 | >167 |
| mIgA1 | 0.04 | 0.93 | >156 | 5.38 | >156 |
| dIgA1 | 0.02 | 0.16 | >74 | 1.90 | >74 |
| sIgA1 | 0.03 | 0.11 | 24.56 | 0.45 | >61 |

IC50 (nM): 0-0.5, 0.5-2, 2-5, >5

| Fold change to IgG | G614 | Alpha | Beta | Delta | Omicron BA.1 |
|---|---|---|---|---|---|
| mIgA1 | 4.41 | 11.22 | – | 1.60 | – |
| dIgA1 | 7.45 | 66.39 | – | 4.53 | – |
| sIgA1 | 6.04 | 91.40 | >6.80 | 19.18 | – |

Fold change: <1, 1-5, 5-10, 10-50, >50

### B rmAb23

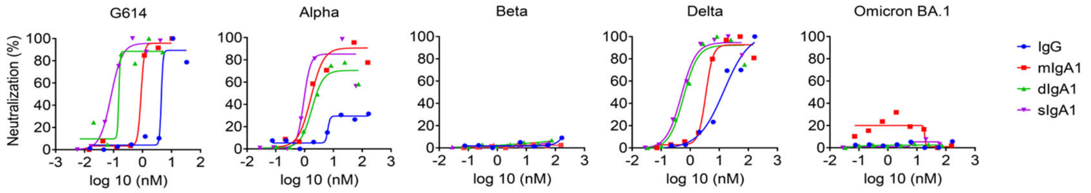


| Neutralization IC50 (nM) | G614 | Alpha | Beta | Delta | Omicron BA.1 |
|---|---|---|---|---|---|
| IgG | 4.44 | >167 | >167 | 14.45 | >167 |
| mIgA1 | 0.89 | 1.66 | >156 | 3.42 | >156 |
| dIgA1 | 0.15 | 1.79 | >74 | 0.59 | >74 |
| sIgA1 | 0.08 | 0.93 | >61 | 0.49 | >61 |

IC50 (nM): 0-0.5, 0.5-2, 2-5, >5

| Fold change to IgG | G614 | Alpha | Beta | Delta | Omicron BA.1 |
|---|---|---|---|---|---|
| mIgA1 | 4.99 | >100.6 | – | 4.23 | – |
| dIgA1 | 29.21 | >93.56 | – | 24.47 | – |
| sIgA1 | 54.44 | >178.82 | – | 29.35 | – |

Fold change: <1, 1-5, 5-10, 10-50, >50

## C XG014

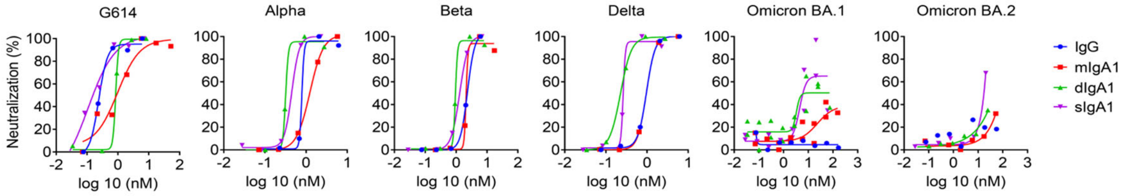


| Neutralization IC50 (nM) | G614 | Alpha | Beta | Delta | Omicron BA.1 | Omicron BA.2 |
|---|---|---|---|---|---|---|
| IgG | 0.24 | 0.79 | 2.34 | 0.97 | >167 | >167 |
| mIgA1 | 0.98 | 1.24 | 2.10 | 0.97 | 23.32 | >156 |
| dIgA1 | 0.85 | 0.32 | 0.95 | 0.23 | 3.29 | >75 |
| sIgA1 | 0.11 | 0.45 | 1.31 | 0.26 | 4.44 | >61 |

IC50 (nM): 0-0.5, 0.5-2, 2-5, >5

| Fold change to IgG | G614 | Alpha | Beta | Delta | Omicron BA.1 | Omicron BA.2 |
|---|---|---|---|---|---|---|
| mIgA1 | 0.24 | 0.64 | 1.11 | 1.01 | >7.16 | – |
| dIgA1 | 0.28 | 2.48 | 2.47 | 4.28 | >50.84 | – |
| sIgA1 | 2.21 | 1.76 | 1.78 | 3.69 | >37.58 | – |

Fold change: <1, 1-5, 5-10, 10-50, >50

**Fig. S6. Dimeric and secretory IgA1 enhanced neutralization activity against variants of concern (VOCs).** (**A** to **C**) Neutralization activity of IgG and IgA1 antibodies 01A05 (A), rmAb23 (B), and XG014 (C) against G614 and VOCs (Alpha, Beta, Delta and Omicron) as determined via microneutralization assay. The $IC_{50}$ and fold-change differences between the IgG and IgA antibody forms are indicated.

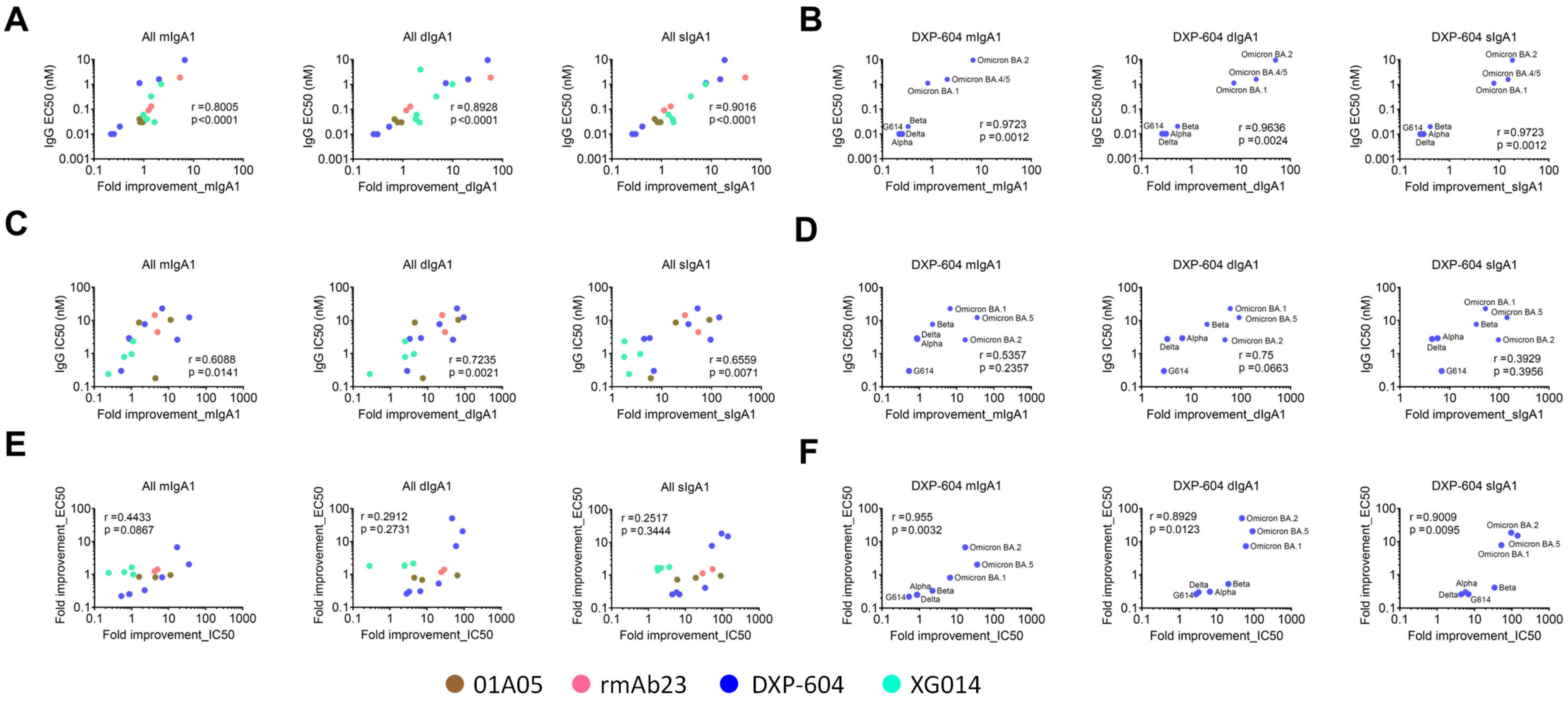


**Fig. S7. Increased neutralization after switching to IgA1 and dimerization was associated with increased RBD binding. (A to D)** Correlation between $EC_{50}$ or $IC_{50}$ of all four IgG antibodies (01A05, rmAb23, DXP-604, and XG014) (A and D) or DXP-604 only (B and D) with increased binding fold changes (A and D) or neutralization fold changes (C and D) after conversion to monomeric (mIgA1), dimeric (dIgA1) or secretory IgA1 (sIgA1). (**E** and **F**) Correlation between the fold change increase in RBD binding and neutralization activity for all four IgG antibodies (E) or DXP-604 only (F) after conversion to mIgA1, dIgA1 and sIgA1.

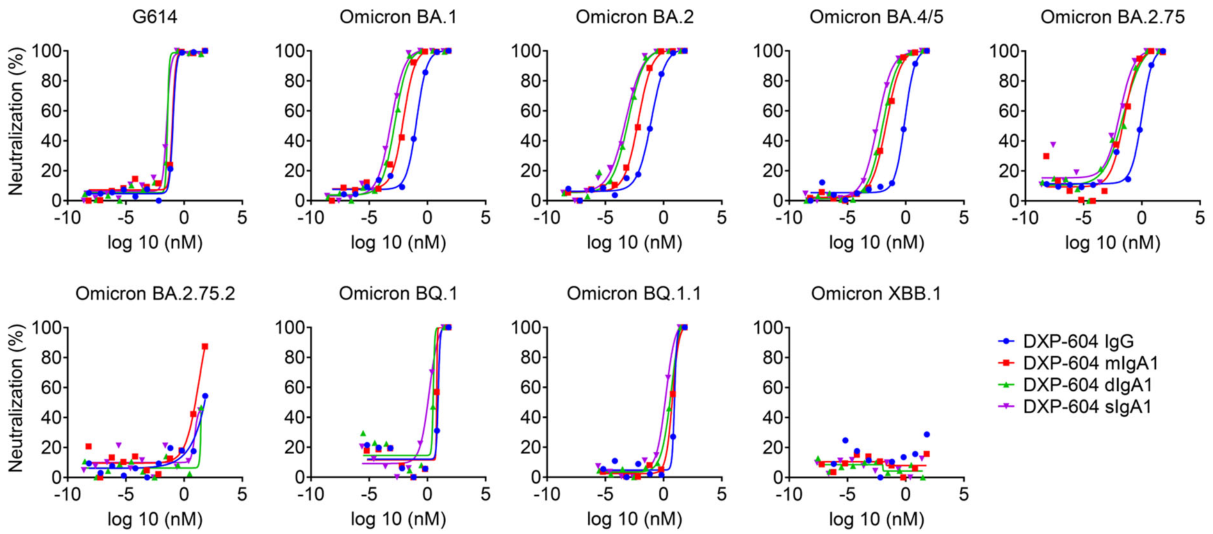


Pseudovirus neutralization IC50 (nM)

| | G614 | Omicron BA.1 | Omicron BA.2 | Omicron BA.4/5 | Omicron BA.2.75 | Omicron BA.2.75.2 | Omicron BQ.1 | Omicron BQ.1.1 | Omicron XBB.1 |
|---|---|---|---|---|---|---|---|---|---|
| IgG | 0.1129 | 0.1205 | 0.0909 | 0.8355 | 1.0160 | >66.67 | 8.4500 | 8.7530 | >66.67 |
| mIgA1 | 0.1007 | 0.0089 | 0.0069 | 0.0221 | 0.0304 | >62.5 | 6.2280 | 5.7670 | >62.5 |
| dIgA1 | 0.0374 | 0.0017 | 0.0009 | 0.0110 | 0.0275 | >29.85 | 3.2010 | 4.8770 | >29.85 |
| sIgA1 | 0.0327 | 0.0008 | 0.0006 | 0.0038 | 0.0130 | >24.39 | 1.4130 | 1.5910 | >24.39 |

IC50 (nM)
<0.001
0.001-0.0
0.01-0.1
>0.1

Fold change to IgG

| | G614 | Omicron BA.1 | Omicron BA.2 | Omicron BA.4/5 | Omicron BA.2.75 | Omicron BA.2.75.2 | Omicron BQ.1 | Omicron BQ.1.1 | Omicron XBB.1 |
|---|---|---|---|---|---|---|---|---|---|
| mIgA1 | 1.12 | 13.54 | 13.17 | 37.81 | 33.43 | – | 1.36 | 1.52 | – |
| dIgA1 | 3.02 | 70.88 | 101.00 | 75.95 | 36.97 | – | 2.64 | 1.79 | – |
| sIgA1 | 3.45 | 150.63 | 151.50 | 219.87 | 77.97 | – | 5.98 | 5.50 | – |

Fold change
<1
1-10
10-50
50-100
>100

**Fig. S8. DXP-604 dimeric and secretory IgA1 enhanced neutralization activity against emerging Omicron subvariants.** DXP-604 neutralization against SARS-CoV-2 S pseudotyped HIV-1-based viruses, including G614 and Omicron BA.1, BA.2, BA.4/5, and recently emerged subvariants (BA.2.75, BA.2.75.2, BQ.1, BQ.1.1 and XBB.1). The $IC_{50}$ and fold-change differences between the IgG and IgA antibody forms are indicated.

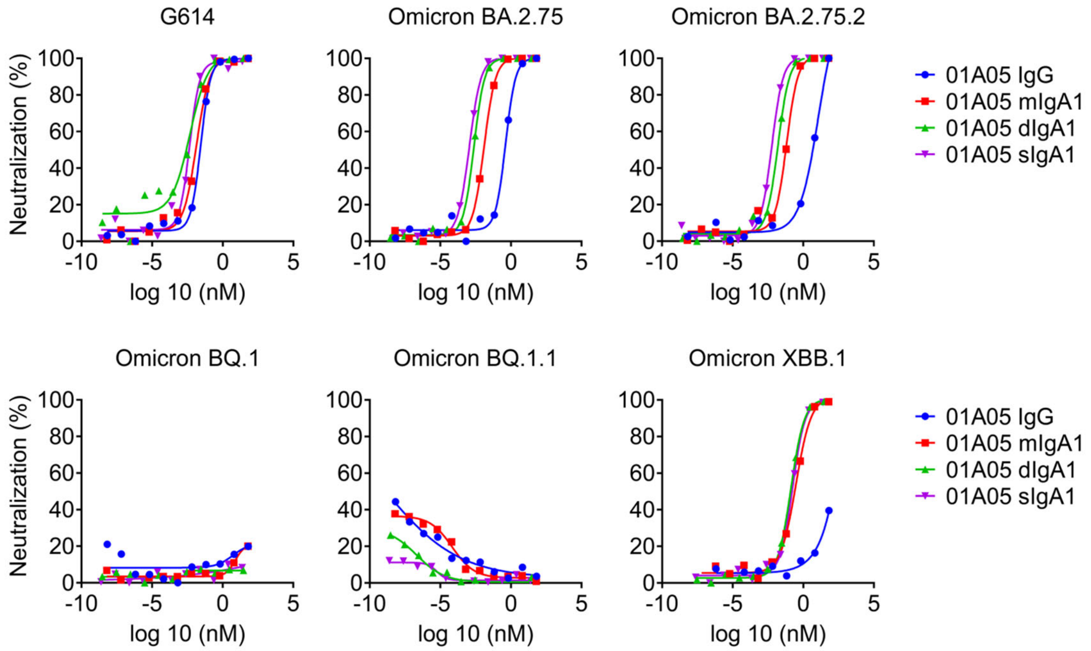


**Pseudovirus neutralization IC50 (nM)**

| | G614 | Omicron BA.2.75 | Omicron BA.2.75.2 | Omicron BQ.1 | Omicron BQ.1.1 | Omicron XBB.1 |
|---|---|---|---|---|---|---|
| IgG | 0.0280 | 0.4195 | 12.0800 | >66.67 | >66.67 | >66.67 |
| mIgA1 | 0.0140 | 0.0127 | 0.0671 | >62.5 | >62.5 | 0.3000 |
| dIgA1 | 0.0048 | 0.0024 | 0.0165 | >29.85 | >29.85 | 0.1347 |
| sIgA1 | 0.0045 | 0.0012 | 0.0060 | >24.39 | >24.39 | 0.1754 |

IC50 (nM)
<0.001
0.001-0.01
0.01-0.1
>0.1

**Fold change to IgG**

| | G614 | Omicron BA.2.75 | Omicron BA.2.75.2 | Omicron BQ.1 | Omicron BQ.1.1 | Omicron XBB.1 |
|---|---|---|---|---|---|---|
| mIgA1 | 2.00 | 32.98 | 180.08 | – | – | >222.23 |
| dIgA1 | 5.85 | 172.99 | 733.01 | – | – | >494.95 |
| sIgA1 | 6.19 | 364.78 | 2023.45 | – | – | >380.10 |

Fold change
<1
1-10
10-50
50-100
>100

**Fig. S9. 01A05 dimeric and secretory IgA1 enhanced neutralization activity against emerging Omicron subvariants.** 01A05 neutralization against SARS-CoV-2 S pseudotyped HIV-1-based viruses, including G614 and recently emerged subvariants (BA.2.75, BA.2.75.2, BQ.1, BQ.1.1 and XBB.1). The $IC_{50}$ and fold-change differences between the IgG and IgA antibody forms are indicated.

**Table S1. Demographic data of vaccinated individuals.**

| Groups | Number | Male/Female | Median age (IQR)[a], years | Median sampling day after vaccination (IQR) |
|---|---|---|---|---|
| Before vaccination | 7 | 3/4 | 37 (28-38) | |
| Inactivated vaccine | | | | |
| 2nd dose | 5 | 2/3 | 29 (28-32) | 35 (16-70) |
| 3rd dose | 6 | 2/4 | 27.5 (26-35) | 68.5 (38-91) |
| mRNA vaccine | | | | |
| 1st dose | 18 | 7/11 | 35.5 (27-40) | 17.5 (14-21) |
| 2nd dose | 36 | 15/21 | 38.5 (28-51) | 28 (17-52) |
| 3rd dose | 22 | 6/16 | 30 (29-44) | 69 (19-126) |
| Heterologous vaccine: Inactivated + 1 dose mRNA vaccine | 13 | 5/8 | 28 (27-30) | 27 (19-36) |
| Infected + mRNA vaccine | 10 | 6/4 | 43 (32-52) | 30 (17-37) |
| Breakthrough infection[b] | 22 | 6/16 | 30.5 (27-45) | 38 (18-138) |

[a] IQR: Interquartile range.

[b] After two or three doses of either inactivated or mRNA vaccine or a combination of both during the Omicron BA.1 wave.

**Table S2. Comparison of DPX604-IgG and IgA1 neutralization activity with commercially available antibodies and antibodies described in the literature.**

| **Antibodies** | **Pseudovirus (pM)** | | | | | | | **Authentic virus (nM)** | | | | | | | | |
|---|---|---|---|---|---|---|---|---|---|---|---|---|---|---|---|---|
| | **G614 or WT** | **Alpha** | **Beta** | **Delta** | **Omicron BA.1** | **Omicron BA.2** | **Omicron BA.4/5** | **G614** | **Alpha** | **Beta** | **Delta** | **Omicron BA.1** | **Omicron BA.2** | **Omicron BA.4** | **Omicron BA.5** | **References[a]** |
| **DXP-604 IgG** | 112.9 | ND[b] | ND | ND | 120.5 | 90.9 | 835.5 | 0.30 | 2.93 | 7.71 | 2.78 | 22.93 | 2.63 | ND | 12.36 | This paper |
| **DXP -604 mIgA1** | 100.7 | ND | ND | ND | 8.90 | 6.9 | 22.1 | 0.56 | 3.41 | 3.42 | 3.15 | 3.43 | 0.16 | ND | 0.35 | This paper |
| **DXP -604 dIgA1** | 37.4 | ND | ND | ND | 1.70 | 0.9 | 11.0 | 0.11 | 0.44 | 0.37 | 0.84 | 0.38 | 0.06 | ND | 0.13 | This paper |
| **DXP -604 sIgA1** | 32.7 | ND | ND | ND | 0.8 | 0.6 | 3.8 | 0.04 | 0.51 | 0.22 | 0.63 | 0.44 | 0.03 | ND | 0.09 | This paper |
| LY-CoV1404 (bebtelovimab, Eli Lilly) | 4 | ND | 14 | ND | 4 | 6 | 6 | 0.070 | 0.027 | 0.047 | 0.053 | 0.107 | ND | 0.08 | 0.10 | Cao et al. 2022 Westendorf et al. 2022. Turelli et al. 2022 |
| COV2-2130 | 16.67 | ND | ND | ND | 20 050 | 42 | 153 | ND | ND | ND | ND | ND | ND | ND | ND | Cao et al. 2022 |
| SA58(BD55-5840) | 6 | ND | ND | ND | 29.33 | 80 | 26 | ND | ND | ND | ND | ND | ND | ND | ND | Cao et al. 2022 |
| SA55(BD55-5514) | 73.3 | ND | ND | ND | 11.33 | 48 | 33.3 | ND | ND | ND | ND | ND | ND | ND | ND | Cao et al. 2022 |
| REGN10987 (Regeneron) | 38 | >66 670 | 13.33 | 33.33 | > 66 670 | 3 930 | 3 470 | ND | ND | ND | ND | ND | ND | ND | ND | Cao et al 2022 |
| S309 | 493.33 | ND | ND | ND | 2 410 | 6 120 | 5 280 | 1.23 | ND | ND | ND | 3.01 | 39.23 | ND | ND | Cao et al. 2022. Case et al. 2022 |

| | | | | | | | | | | | | | | | | |
|---|---|---|---|---|---|---|---|---|---|---|---|---|---|---|---|---|
| REGN10933+ REGN10987 (Regeneron) | 33.33 | ND | ND | ND | >66 670 | 5 470 | 4 730 | | ND | ND | ND | ND | ND | ND | ND | Cao et al. 2022 |
| AZD1061 (Astra Zeneca) | 13.33 | ND | ND | ND | 2 050 | 53.33 | 100 | 0.197 | ND | ND | ND | 40.52 | 0.21 | 0.81 | 0.95 | Case et al. 2022 Tuekprakhon et al. 2022. Turelli et al. 2022 |
| AZD7442 (Astra Zeneca) (Tixagevimab+ Cilgavimab) | 6.67 | ND | ND | ND | 1 550 | 53.33 | 433 | 0.043 | ND | ND | ND | 1.11 | 0.24 | ND | ND | Case et al 2022 Tuekprakhon et al. 2022 |

[a] Cao, Y. et al. BA.2.12.1. BA.4 and BA.5 escape antibodies elicited by Omicron infection. *Nature* **608,** 593–602 (2022).

Case, J.B. et al. Resilience of S309 and AZD7442 monoclonal antibody treatments against infection by SARS-CoV-2 Omicron lineage strains. *Nat. Commun.* **13,** 3824 (2022).

Tuekprakhon, A. et al. Antibody escape of SARS-CoV-2 Omicron BA.4 and BA.5 from vaccine and BA.1 serum. *Cell* **185,** 2422-2433.e13 (2022).

Westendorf, K. et al. LY-CoV1404 (bebtelovimab) potently neutralizes SARS-CoV-2 variants. *Cell rep.* **39,** 110812 (2022).

Turelli, P. et al. P2G3 human monoclonal antibody neutralizes SARS-CoV-2 Omicron subvariants including BA.4 and BA.5 and Bebtelovimab escape mutants. bioRxiv (2022), https://doi.org/10.1101/2022.07.28.501852.
